# Massively parallel single-cell chromatin landscapes of human immune cell development and intratumoral T cell exhaustion

**DOI:** 10.1101/610550

**Authors:** Ansuman T. Satpathy, Jeffrey M. Granja, Kathryn E. Yost, Yanyan Qi, Francesca Meschi, Geoffrey P. McDermott, Brett N. Olsen, Maxwell R. Mumbach, Sarah E. Pierce, M. Ryan Corces, Preyas Shah, Jason C. Bell, Darisha Jhutty, Corey M. Nemec, Jean Wang, Li Wang, Yifeng Yin, Paul G. Giresi, Anne Lynn S. Chang, Grace X.Y. Zheng, William J. Greenleaf, Howard Y. Chang

**Affiliations:** Center for Personal Dynamic Regulomes, Stanford University School of Medicine, Stanford, CA 94305, USA; Department of Pathology, Stanford University School of Medicine, Stanford, CA 94305, USA; Department of Genetics, Stanford University School of Medicine, Stanford, CA 94305, USA; Biophysics Program, Stanford University School of Medicine, Stanford, CA 94305, USA; Cancer Biology Program, Stanford University School of Medicine, Stanford, CA 94305, USA; Department of Dermatology, Stanford University School of Medicine, Redwood City, CA 94063, USA; 10x Genomics, Inc., Pleasanton, CA 94566, USA; Department of Applied Physics, Stanford University, Stanford, CA 94305, USA; Chan Zuckerberg Biohub, San Francisco, CA 94158, USA; Howard Hughes Medical Institute, Stanford University School of Medicine, Stanford, CA 94305, USA

## Abstract

Understanding complex tissues requires single-cell deconstruction of gene regulation with precision and scale. Here we present a massively parallel droplet-based platform for mapping transposase-accessible chromatin in tens of thousands of single cells per sample (scATAC-seq). We obtain and analyze chromatin profiles of over 200,000 single cells in two primary human systems. In blood, scATAC-seq allows marker-free identification of cell type-specific *cis*- and *trans*-regulatory elements, mapping of disease-associated enhancer activity, and reconstruction of trajectories of differentiation from progenitors to diverse and rare immune cell types. In basal cell carcinoma, scATAC-seq reveals regulatory landscapes of malignant, stromal, and immune cell types in the tumor microenvironment. Moreover, scATAC-seq of serial tumor biopsies before and after PD-1 blockade allows identification of chromatin regulators and differentiation trajectories of therapy-responsive intratumoral T cell subsets, revealing a shared regulatory program driving CD8^+^ T cell exhaustion and CD4^+^ T follicular helper cell development. We anticipate that droplet-based single-cell chromatin accessibility will provide a broadly applicable means of identifying regulatory factors and elements that underlie cell type and function.

## Introduction

Tissues are comprised of a complex collection of cell types, each type specialized to its functional role. Understanding this inherent functional complexity often requires technologies that measure properties of single cells, rather than of the system as a whole. This concept is exemplified in the immune system, where effective responses to a wide array of pathogens or cancer are orchestrated by the sequential activity of more than a hundred specialized cell types. While recent studies have developed a number of robust technologies to measure RNA or protein expression in single cells, technologies for assessing chromatin accessibility in single cells are comparatively lacking.

Cell type-specific gene expression in eukaryotic cells is regulated by millions of *cis*-acting DNA elements (e.g. enhancers and promoters) and thousands of *trans*-acting factors (e.g. transcription factors [TFs] and regulatory RNAs)^1^. Functionally distinct cell types exhibit distinct gene expression programs that are brought about by changes in the activity of *cis*-elements through their interplay with *trans*-factors. We previously developed the assay for transposase-accessible chromatin using sequencing (ATAC-seq), which enables the enumeration of active DNA regulatory elements by direct transposition of sequencing adapters into accessible chromatin with the hyperactive transposase Tn5^2^. This method can reveal several layers of gene regulation in a single assay, including the genome-wide identification of *cis*-elements, inference of TF binding and activity, and nucleosome positions^2–4^. Importantly, ATAC-seq is applicable to low-cell number samples^5^, and even single cells^6,7^, which has enabled epigenomic profiling of primary samples with newfound precision. In studies to date, single-cell ATAC-seq (scATAC-seq) has been used to map cell-to-cell variability and rare single-cell epigenomic phenotypes across diverse biological processes, including in healthy and malignant immune cells^8–12^. However, the widespread adoption of this technique has been hindered by the difficulty and cost of performing the assay at scale while maintaining high data quality.

Here we report a method to perform scATAC-seq in nanoliter-sized droplets, which enables the generation of high-quality single-cell chromatin accessibility profiles at massive scale. To demonstrate the performance and utility of this method, we analyzed primary cells in two biological contexts. First, we mapped the single-cell chromatin accessibility landscape of blood formation in bone marrow and blood samples from healthy humans. This landscape revealed diverse chromatin states of progenitor cells and the regulatory trajectories of their differentiation into effector cell types. Second, we performed scATAC-seq in primary tumor biopsies from patients with basal cell carcinoma (BCC) receiving anti-programmed cell death protein 1 (PD-1) immunotherapy (PD-1 blockade). Single-cell deconvolution of the tumor microenvironment (TME) revealed distinct types of immune, stromal, and malignant cells, and analysis of intratumoral T cells identified epigenetic regulators of therapy-responsive T cell subtypes, including CD8^+^ exhausted (TEx) and CD4^+^ T follicular helper (Tfh) cells. Altogether, we report scATAC-seq profiles of over 200,000 cells, demonstrating that this platform enables the unbiased discovery of cell types and regulatory DNA elements across diverse biological systems.

## Results

### Droplet-based platform for measuring single-cell chromatin accessibility

We developed a method to perform scATAC-seq in droplets using the Chromium platform (10X Genomics) previously employed to measure single-cell transcriptomes^13^ or paired single-cell transcriptomes and T cell- or B cell-receptor sequences^14^ (Fig. 1a, **Supplementary Fig. 1a**). In this scATAC-seq approach, nuclei are first isolated from a single-cell suspension and transposed in bulk with the transposase Tn5. Transposed nuclei are then loaded onto an 8-channel microfluidic chip for Gel bead in Emulsion (GEM) generation. Each gel bead is functionalized with newly-designed single-stranded barcoded oligonucleotides that consists of: (1) a 29 basepair (bp) sequencing adapter, (2) a 16bp barcode selected from ~750,000 designed sequences to index GEMs, and (3) the first 14bp of Read 1N, which serves as the priming sequence in the linear amplification reaction to incorporate barcodes to transposed DNA (**Supplementary Fig. 1a**). In each channel, ~100,000 GEMs are formed per experiment, resulting in the encapsulation of tens of thousands of cells in GEMs per microfluidic chip. Approximately 80% of GEMs contain a single gel bead, and nuclei are loaded at a limiting dilution to minimize co-occurrence of multiple cells in the same GEM. After GEM generation, gel beads are dissolved, and the oligonucleotides are released for linear amplification of transposed DNA. Finally, the emulsion is broken, and barcoded DNA is pooled for PCR amplification to generate indexed libraries, which are compatible with high-throughput sequencing. In the sequencing reaction, reads 1N and 2N contain the DNA insert, while the index reads, i5 and i7, capture the cell barcodes and sample indices, respectively.

**Figure 1.**
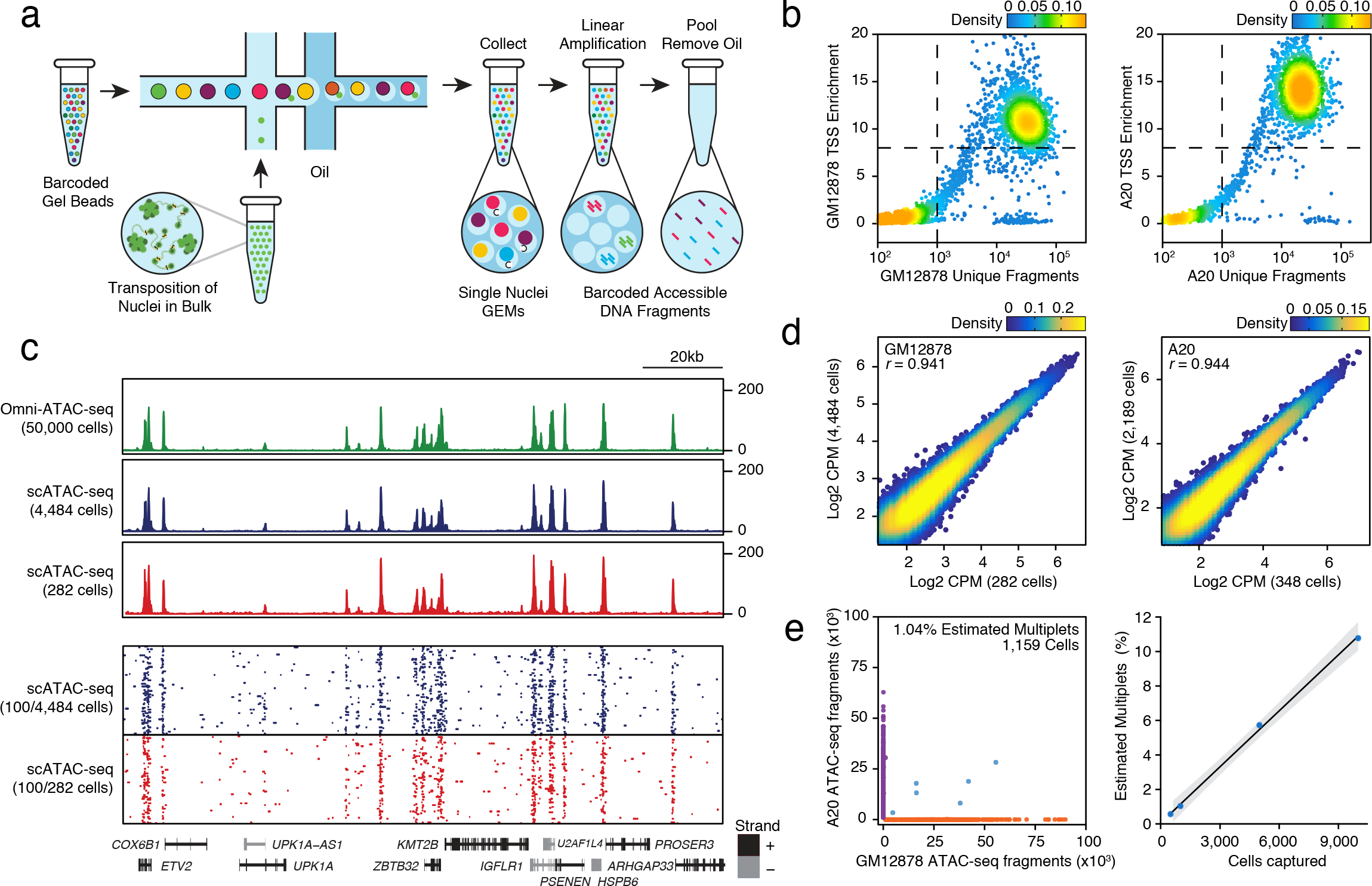
Massively parallel scATAC-seq in droplets. (**a**) Schematic of scATAC-seq in droplets. (**b**) ATAC-seq data quality control filters in human (GM12878) and mouse (A20) B cells at 5,000 cell loading. Shown are the number of unique ATAC-seq nuclear fragments in each single cell (each dot) compared to TSS enrichment of all fragments in that cell. Dashed lines represent the filters for high-quality single-cell data (1,000 unique nuclear fragmentts and TSS score greater than or equal to 8). Data is representative of 4 independent experiments. (**c**) Genome tracks showing the comparison of aggregate scATAC-seq profiles to bulk Omni-ATAC-seq profiles from GM12878 B lymphoblasts (top panel). scATAC-seq profiles were obtained from two independent mixing experiments, in which either 4,484 (from 10,000 cell loading) or 282 (from 500 cell loading) cells were assayed, as indicated. The bottom panel shows accessibility profiles of 100 random single GM12878 cells from each experiment. Each pixel represents a 100bp region. (**d**) One to one plots of log-normalized reads in ATAC-seq peaks in aggregate scATAC-seq profiles. Aggregate profiles in GM12878 (left) and A20 (right) cells are derived from two individual mixing experiments as in b, in which the indicated number of cells were assayed. ATAC-seq peaks were identified in Omni-ATAC-seq profiles from 50,000 cells^5^. (**e**) Human (GM12878)/mouse (A20) cell mixing experiment showing proportion of single-cell libraries with both mouse and human ATAC-seq fragments (left). The right panel shows proportion of mouse/human multiplets detected when cell loading concentration was varied across four independent experiments. Shaded lines indicate 95% confidence interval.

To assess the performance of this method, we generated scATAC-seq libraries from species-mixing experiments, in which we pooled human (GM12878) and mouse (A20) B cell nuclei. The nuclei from each cell type were first transposed in bulk, and then an even mixture of each cell type was loaded at various loadings onto microfluidic chips. The resulting libraries were sequenced and processed through a single-cell analysis pipeline to de-multiplex reads, assign cell barcodes, align fragments to the human or mouse reference genome, and de-duplicate fragments generated by PCR amplification (Cell Ranger ATAC; **Methods**). We first evaluated the quantity and quality of all scATAC-seq data, regardless of species of origin, and used previously described cut-offs of 1,000 unique nuclear fragments per cell and a transcription start site (TSS) enrichment score of 8 to exclude low-quality cells from further downstream analysis (**Methods**)^15^. Single cells passing filter yielded on average 23.24 × 10^3^ unique fragments mapping to the nuclear genome, and approximately 40.5% of Tn5 insertions were within peaks present in aggregated ATAC-seq profiles from all cells, comparable to previously published high-quality scATAC-seq and bulk ATAC-seq profiles (Fig. 1b-c **and Supplementary Fig. 1b**)^6,10,15^. Accordingly, scATAC-seq profiles exhibited fragment size periodicity and a high enrichment of fragments at TSSs, and aggregate profiles from multiple independent experiments were highly correlated (Fig. 1d **and Supplementary Fig. 1c**). Finally, analysis of mouse and human fragments in single cells confirmed a low-rate of estimated multiplets (12/1,159 cells, ~1%; Fig. 1e). A cell titration experiment with four different cell loading concentrations showed a linear relationship between the observed multiplet rate and the number of recovered cells (~1% multiplets per 1,000 cells recovered), consistent with Poisson loading of nuclei and similar observations for single-cell transcriptomes in droplet platforms (Fig. 1e)^13,16^. Therefore, to minimize potential multiplets, we typically aimed to capture <6,000 nuclei per channel.

### Validation of rare cell detection and performance in complex archival samples

We next sub-sampled scATAC-seq data *in silico* to assess the technical performance of scATAC-seq with different data quantity (fragments/cell) and cell number (**Supplementary Fig. 1d-e**). A comparison of each sub-sampled profile to the aggregate profile from all data demonstrated that a relatively complete list of *cis*-elements could be identified by aggregating ~200 cells and at least 10,000 fragments/cell. Aggregate profiles from ~200 cells could achieve the confident discovery of ~80% of ATAC-seq peaks from total profiles, and an overall Pearson correlation of *r*~0.9 for all reads in peaks (**Supplementary Fig. 1d**). With this information in mind, we devised a data analysis workflow for peak calling and clustering (**Supplementary Fig. 1f, Methods**). In this workflow, single-cell libraries were first processed with Cell Ranger and filtered. Next, we performed an ‘initial’ clustering analysis by partitioning the genome into 2.5 kb windows and counting Tn5 insertions in each window, as described previously^7,9^. We then performed single-cell Latent Semantic Indexing (LSI) and clustered cells using Shared Nearest Neighbor (SNN) clustering (SNN; Seurat^17^) with the top 20,000 accessible windows, requiring that each cluster contain at least 200 cells. These ‘initial’ clusters were then used to identify confident ATAC-seq peaks (using MACS2^18^) in single-cell clusters and to generate a merged peak set that represented the full epigenetic diversity of all cells. Finally, a cell-by-peak counts matrix was created and used for ‘final’ single-cell clustering and downstream analysis, in which each cluster could contain any number of cells (**Methods**).

We tested this analysis approach with two quality-control experiments. First, we generated a series of synthetic cell mixtures, in which human monocytes were isolated from peripheral blood mononuclear cells (PBMCs) and mixed with sorted human T lymphocytes in various ratios spanning a 1,000-fold detection range (**Supplementary Fig. 2a-b, Supplementary Table 1**). We then performed scATAC-seq and determined whether we could resolve each population in an unsupervised analysis. As expected, scATAC-seq analysis of 50:50 mixtures identified two distinct populations of cells, which demonstrated high accessibility of open chromatin regions linked either to monocyte-specific genes (i.e. *CD14*, *CSF1R*, *TREML4*), or to T cell-specific genes (i.e. *CD3E*, *CD4*, *CD8A*; **Supplementary Fig. 2a and Methods**). Importantly, this analysis could also resolve small populations, which represented either 1/100 or 1/1,000 of total cells, demonstrating that even rare cell-types could be identified in this manner (**Supplementary Fig. 2b**). Second, we compared the performance of scATAC-seq analysis in fresh versus frozen PBMCs (**Supplementary Fig. 2c-f**). We isolated nuclei from either fresh PBMCs, viably-frozen PBMCs, or viably-frozen PBMCs that were sorted for live cells and performed scATAC-seq (**Methods**). As seen in previous datasets, we confirmed that scATAC-seq profiles passing filter yielded approximately the same quantity and quality of ATAC-seq data, regardless of sample origin, and that aggregate profiles from fresh and frozen cells were highly correlated (**Supplementary Fig. 2c-d**)^11^. Importantly, cell type-specific clusters from each PBMC sample clustered together, and frozen samples recapitulated the majority of ATAC-seq peaks discovered in fresh samples (AUC: 0.809; **Supplementary Fig. 2e-f**). Altogether, these results quantify the high similarity of scATAC-seq data generated in different batches and across different sample preparation conditions and demonstrate the ability to discover rare single cells in heterogeneous mixtures.

**Figure 2.**
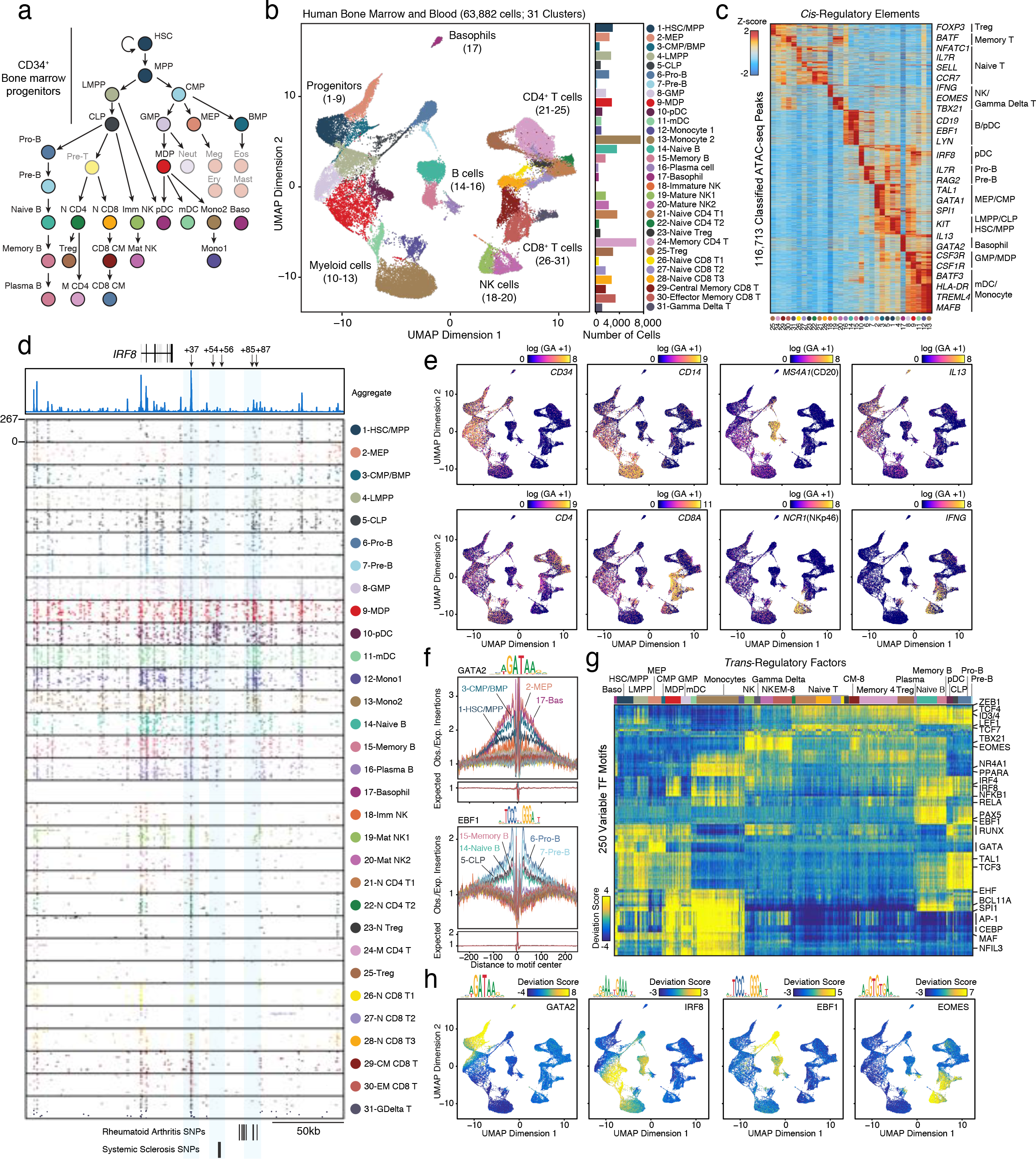
Single-cell chromatin accessibility of human hematopoiesis. (**a**) Schematic of progenitor and end-stage cell types in human hematopoiesis. (**b**) UMAP projection of 63,882 scATAC-seq profiles of bone marrow and peripheral blood immune cell types. Dots represent individual cells, and colors indicate cluster identity (labeled on the right). Bar plot indicates the number of scATAC-seq profiles in each cluster of cells. (**c**) Heatmap of Z-scores of 116,713 *cis*-regulatory elements in scATAC-seq clusters derived from (b). Gene labels indicate the nearest gene to each regulatory element. (**d**) Single-cell chromatin accessibility in the *IRF8* locus. Each box shows scATAC-seq profiles from 100 representative single cells from each cluster. Each pixel represents a 200bp region. The top genome track shows the aggregate accessibility profile from all cells combined. (**e**) UMAP projection colored by log-normalized gene activity scores demonstrating the accessibility of *cis*-regulatory elements linked (computed from linked accessibility of distal peaks to peaks at gene promoters using Cicero) to the indicated gene. For example, the top left plot demonstrates the accessibility score for *cis*-elements linked to the promoter of the hematopoietic progenitor gene *CD34*. (**f**) Example TF footprints of GATA2 and EBF1 with motifs in the indicated scATAC-seq clusters. The Tn5 insertion bias track is shown below. (**g**) Heatmap representation of ATAC-seq chromVAR bias-corrected deviations in the 250 most variable TFs across all scATAC-seq clusters. Single-cell cluster identities are indicated at the top of the plot. (**h**) UMAP projection of scATAC-seq profiles colored by chromVAR TF motif bias-corrected deviations for the indicated factors.

### Unbiased single-cell accessible chromatin landscape of human hematopoiesis

To further demonstrate this method in primary samples, we performed experiments in human immune cells. The immune system relies on the continuous differentiation of hematopoietic stem cells (HSCs) into functionally-specialized cell types, a process which is tightly controlled by the expression of cell-type-specific genes and maintained by epigenetic programs (Fig. 2a). Since this system has been extensively studied using single-cell methods, we reasoned that it might be an ideal system to validate the interpretation of scATAC-seq data at scale. We generated scATAC-seq libraries from peripheral blood (PB) and bone marrow (BM) cells from 16 healthy individuals and sampled cells both in an unbiased fashion, analyzing total PB and BM cells, or after cell sorting to enrich for certain cell phenotypes (**Supplementary Fig. 3a and Supplementary Table 2**). The purpose of this sampling strategy was two-fold: (1) to validate scATAC-seq-defined cluster identities by standard cell surface marker phenotypes, and (2) to uncover additional heterogeneity within surface-marker defined populations with single-cell measurements. In total, we generated high-quality scATAC-seq profiles from 61,806 cells, and cells passing filter yielded on average 15.6 × 10^3^ fragments mapping to the nuclear genome, and approximately 40.5% of Tn5 insertions were within peaks identified in the aggregate ATAC-seq profile from all cells (**Supplementary Fig. 3b-c**). Importantly, the quality of scATAC-seq profiles was highly uniform across individuals, samples, and cell types, and the number of fragments per cell and TSS enrichment scores were on par with high-quality scATAC-seq datasets in primary immune cells generated with other technologies (**Supplementary Fig. 3d-e**)^11,12^.

**Figure 3.**
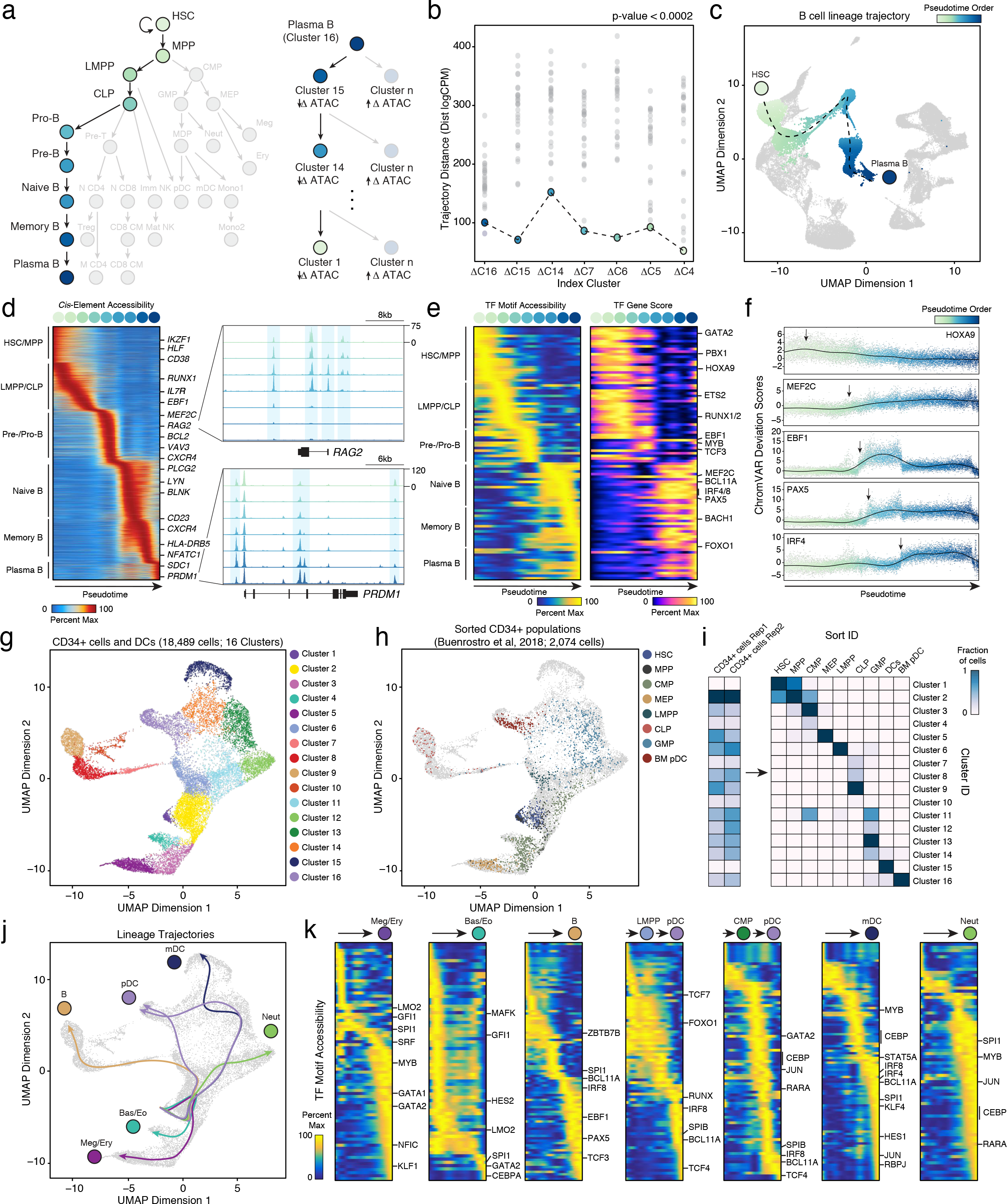
Epigenomic differentiation trajectories of human immune cell types. (**a**) Differentiation trajectory of HSCs to terminal plasma B cells (left). Reverse reconstruction of B cell differentiation trajectory using scATAC-seq profiles (right; see Methods). Differences between the aggregate plasma B cell scATAC-seq profile and all other clusters are calculated. Trajectory is tested against a nearest neighbor approach; the cluster with the most similarity (lowest trajectory distance) to the cluster of interest is identified as the immediate precursor cluster. (**b**) Trajectory distance calculations for the terminal plasma B cell cluster (Cluster 16). Dots represent comparisons between the cluster of interest (labeled at the bottom) and every cluster that has not previously identified. P-value < 0.0002 represents 5000 random simulations of trajectory ordering. Pseudotime representation of plasma B cell differentiation from HSCs. The dashed line represents double-spline fitted trajectory across pseudotime. (**d**) Pseudotime heatmap ordering of the top 10,000 variable *cis*-regulatory elements across B cell differentiation (left). Zoom-in genome tracks show representation behavior of *cis*- elements accessible early (top) and late in B cell differentiation (bottom). (**e**) Pseudotime heatmap ordering of chromVAR TF motif bias-corrected deviations across B cell differentiation (left). TF motifs are filtered for genes that are highly active (defined as the average percentile between total gene activity and variability) that also demonstrate similarly dynamic gene scores across differentiation (R > 0.35 and FDR < 0.001). Heatmap of TF gene scores is shown on the right. (**f**) chromVAR bias-corrected deviation scores for the indicated TFs across B cell pseudotime. Each dot represents the deviation score in an individual pseudotime-ordered scATAC-seq profile. The line represents the smoothed fit across pseudotime and chromVAR deviation scores. (**g**) Sub-clustering UMAP projection of 18,489 CD34^+^ BM progenitors and DCs (Clusters 1-6 and 8-11 from full hematopoiesis). scATAC-seq profiles are colored by cluster identity, as labeled on the right. (**h**) UMAP projection of progenitor populations, highlighted are the sorted progenitor populations from Buenrostro et al., 2018^11^. Greyed out are the cells assayed in this study. (**i**) Confusion matrix of sorted progenitor populations showing the proportion of each population in clusters defined in (g). (**j**) Lineage trajectories for the indicated cell types, calculated as described in (a). Lines represent double-spline fitted trajectories across pseudotime. (**k**) Pseudotime heatmap ordering of chromVAR TF motif bias-corrected deviations in the indicated lineage trajectory. TF motifs are filtered for genes as described in e.

We clustered scATAC-seq profiles with LSI followed by SNN clustering and visualized clusters with uniform manifold approximation and projection (UMAP), a nonlinear dimensionality-reduction technique that preserves local and aspects of global inter-cluster relationships^19^. We identified 31 scATAC-seq clusters, which we classified using three parallel approaches: (1) chromatin accessibility of individual *cis*-elements (ATAC-seq peaks), (2) gene activity scores, which were computed from the accessibility of several enhancers linked to a single gene promoter^20^, and (3) TF activity, as computed from the accessibility of thousands of TF binding sites genome-wide in each single cell^4^. All three approaches are based on a ‘bottom-up’ analysis of scATAC-seq data and do not require prior knowledge from RNA-seq datasets or reference bulk ATAC-seq profiles.

Using the first approach, we identified a total of 571,400 equal-width non-overlapping *cis*-elements across all scATAC-seq clusters, and approximately 20.4% of elements (116,713) exhibited significant cell-type specific accessibility in a single cluster (mean 6,208 peaks per cluster, FDR < 0.01; **Methods**). Annotation of cell types through the identification of neighboring genes to cluster-specific *cis*-elements demonstrated that scATAC-seq profiles spanned the continuum from early hematopoietic progenitors to end-stage cell types (Fig. 2b-c). For example, Clusters 2-4 demonstrated accessibility at *cis*- elements neighboring key myeloid progenitor genes, including *KIT*, *GATA1*, *TAL1*, and *SPI1*, while Clusters 14-16 demonstrated accessibility at *cis*-elements neighboring B cell genes, including *CD19*, *EBF1*, and *LYN* (Fig. 2c **and Supplementary Fig. S4a**). Importantly, clustering of scATAC-seq profiles could identify relatively rare cell types, such as basophils, as well as known cell type distinctions and subsets, such as the distinction between CD4^+^ and CD8^+^ T cells, and the presence of phenotypically-distinct T cell subsets, such as regulatory CD4^+^ T cells (Treg; Fig. 2b-c). Moreover, scATAC-seq analysis identified unique cell-type specific regulatory elements even within a single gene locus. For example, although the transcription factor IRF8 is critical for the function of many immune cell types, we observed unique accessibility of the +85kb and +87kb enhancers in the *IRF8* locus in myeloid cells, and of the +54kb and +56kb enhancers in plasmacytoid dendritic cells (pDCs), while the +37kb enhancer was accessible in nearly all immune lineages (Fig. 2d). These findings are in line with previously-identified *Irf8* super-enhancers in dendritic cells^21^, and potentially inform the cellular impact of genetic variants associated with autoimmune disease present in this locus^22^.

Although *cis*-element analysis can be informative, this measurement is naturally sparse in single cells, as it is limited by the DNA copy number (2 alleles per element in a diploid genome). Therefore, in the second analysis approach, we asked whether we could further support cluster identities using gene activity scores (henceforth referred to as ‘gene scores’), which are computed as the normalized aggregate accessibility of several enhancers linked to a single gene promoter^20^. We first identified all enhancer-promoter (E-P) connections genome-wide with Cicero, an algorithm that links pairs of DNA elements based on co-accessibility in scATAC-seq data^20^. This method identified 149,309 total E-P connections across all scATAC-seq clusters, with a median of 6 enhancers linked to each gene promoter (**Methods**). We independently validated E-P connections obtained from this analysis using two orthogonal datasets from primary human immune cells. First, we compared E-P connections identified with Cicero to chromosome conformation signal obtained from H3K27ac HiChIP experiments in T cells^23^ and found that enhancers linked to gene promoters showed significant enrichment for HiChIP enhancer interaction signal (EIS), compared to neighboring genomic regions (**Supplementary Fig. 4b**). Second, we compared Cicero E-P contacts to expression quantitative trait loci (eQTLs; Genotype-Tissue Expression [GTEx] database^24^) and found enrichment of eQTLs in linked contacts, particularly when eQTLs were also identified in immune cells (**Supplementary Fig. 4c; Methods**). Together, these results confirm that globally, E-P contacts identified with this method are supported by three-dimensional genome conformation and by functional perturbations of enhancers.

**Figure 4.**
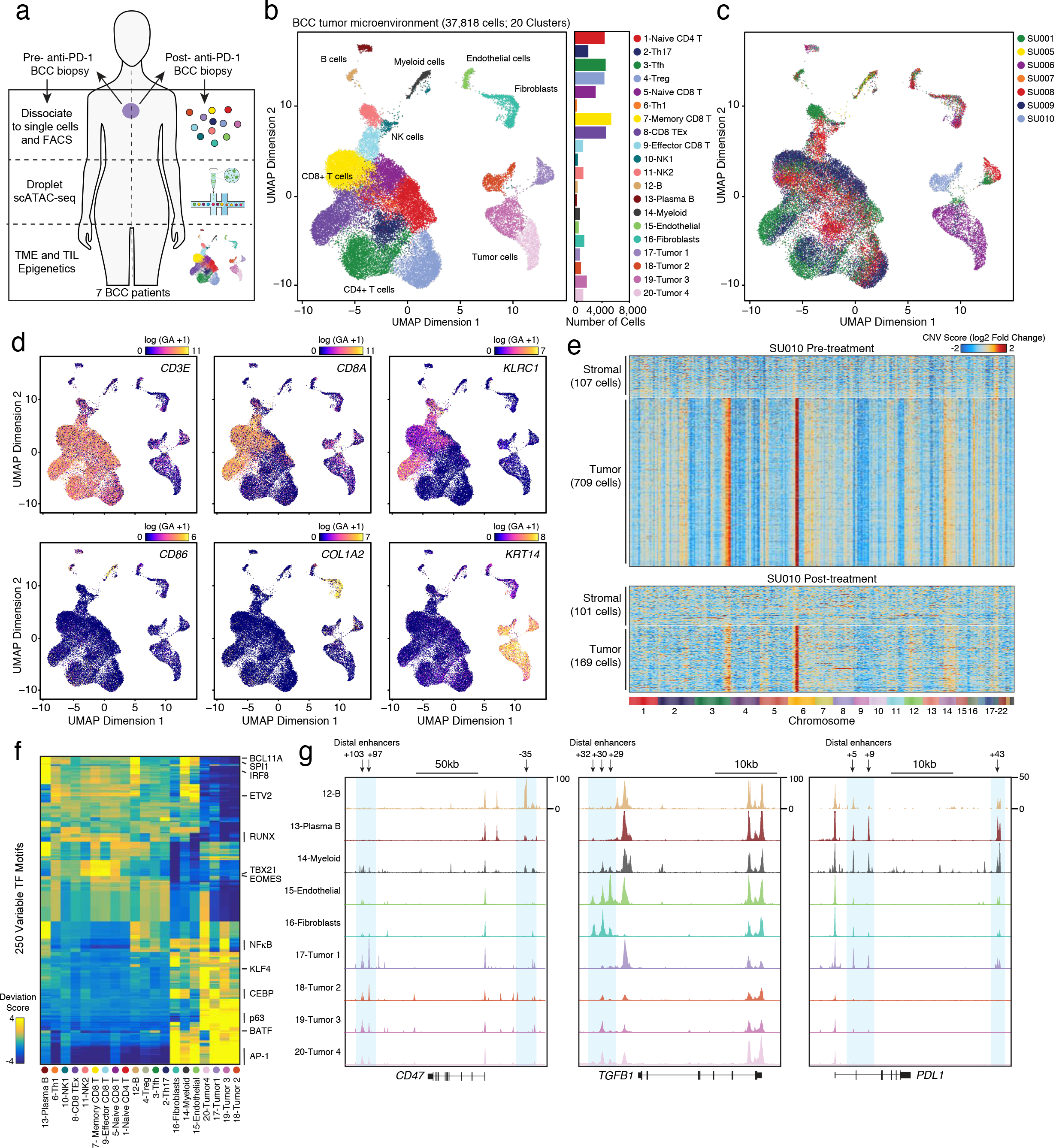
Single-cell regulatory landscape of the BCC TME. (**a**) Schematic of analysis of BCC samples. (**b**) UMAP projection of 37,818 scATAC-seq profiles of BCC TME cell types. Dots represent individual cells, and colors indicate cluster identity (labeled on the right). Bar plot indicates the number of cells in each cluster of cells. (**c**) UMAP projection colored by patient of origin, as indicated on the right. (**d**) UMAP projection colored by log normalized gene activity scores demonstrating the accessibility of *cis*-regulatory elements linked (using Cicero) to the indicated gene. (**e**) Estimated Copy-Number Variation (log2 Fold Change to GC matched background) from scATAC-seq data. Stromal cells include endothelial cells and fibroblasts. (**f**) Heatmap representation of ATAC-seq chromVAR bias-corrected deviations in the 250 most variable TFs across all scATAC-seq clusters. Cluster identities are indicated at the bottom of the plot. (**g**) Genome tracks of aggregate scATAC-seq data, clustered as indicated in (b). Arrows indicate the position and distance (in kb) of distal enhancers in each gene locus.

We next calculated single-cell accessibility at E-P connections for each gene and projected gene scores for immune lineage-defining genes onto scATAC-seq clusters (Fig. 2e **and Methods**). Indeed, we found that gene scores supported prior *cis*-element-defined cluster identities, and that many of these scores were relatively robust in each cell within a cluster. For example, the *CD34* gene score identified hematopoietic progenitors, the *CD14* gene score identified monocyte and myeloid dendritic cells (mDCs), and the *CD20* gene score identified B cells (Fig. 2e **and Supplementary Fig. S4d-e**). Again, this analysis robustly identified rare cell types, for example demonstrating high *IL13* gene scores in basophils, and immune cell subsets, for example identifying high *FOXP3* scores in Tregs (Fig. 4e). Across all single cells, we identified 5,977 gene scores that exhibited scATAC-seq cluster-specific activity, reflecting known and novel markers for each cell type (**Supplementary Fig. 4d**).

Finally, in the third analysis approach, we measured chromatin accessibility at all *cis*-elements sharing a common feature (such as a TF binding motif) using chromVAR^4^. To demonstrate the viability of this method, we analyzed aggregate scATAC-seq cluster profiles for expected differences at binding sites for known cell type-specific TFs. Indeed, genome-wide accessibility at binding sites for GATA2, a lineage-determining factor for megakaryocyte, erythrocyte, and basophil lineages^25^, was increased in megakaryocyte-erythroid progenitors (MEPs), in basophils, and in common myeloid progenitors (CMPs), as expected (Fig. 2f). Similarly, the accessibility of binding sites for EBF1, the lineage-determining factor for B cells^26^, was increased in naïve, memory, and plasma B cells, as well as in early B cell progenitors (Fig. 2f). Since DNA bound by TFs is protected from transposition by Tn5, visualization of each TF profile showed local chromatin accessibility changes surrounding the binding ‘footprint’ (Fig. 2f **and Supplementary Fig. 5a**). Next, we computed the genome-wide accessibility for all TF motifs in each single cell using chromVAR (referred to as ‘chromVAR TF deviation’), which revealed shared and unique regulatory programs across immune cell types (Fig. 2g-h **and Supplementary Fig. 5b**). For example, mDCs and B cells shared activity at sites containing BCL11A, SPI1, and IRF factor motifs, but demonstrated unique activity at sites containing motifs for CEBP factors and EBF1, respectively (Fig. 2g-h). Similarly, TBX21- and EOMES-bound sites were active in both NK and T cell populations; however, only T cells showed accessibility at sites containing motifs for the T cell lineage-determining factor TCF7 (Fig. 2g-h)^27^.

**Figure 5.**
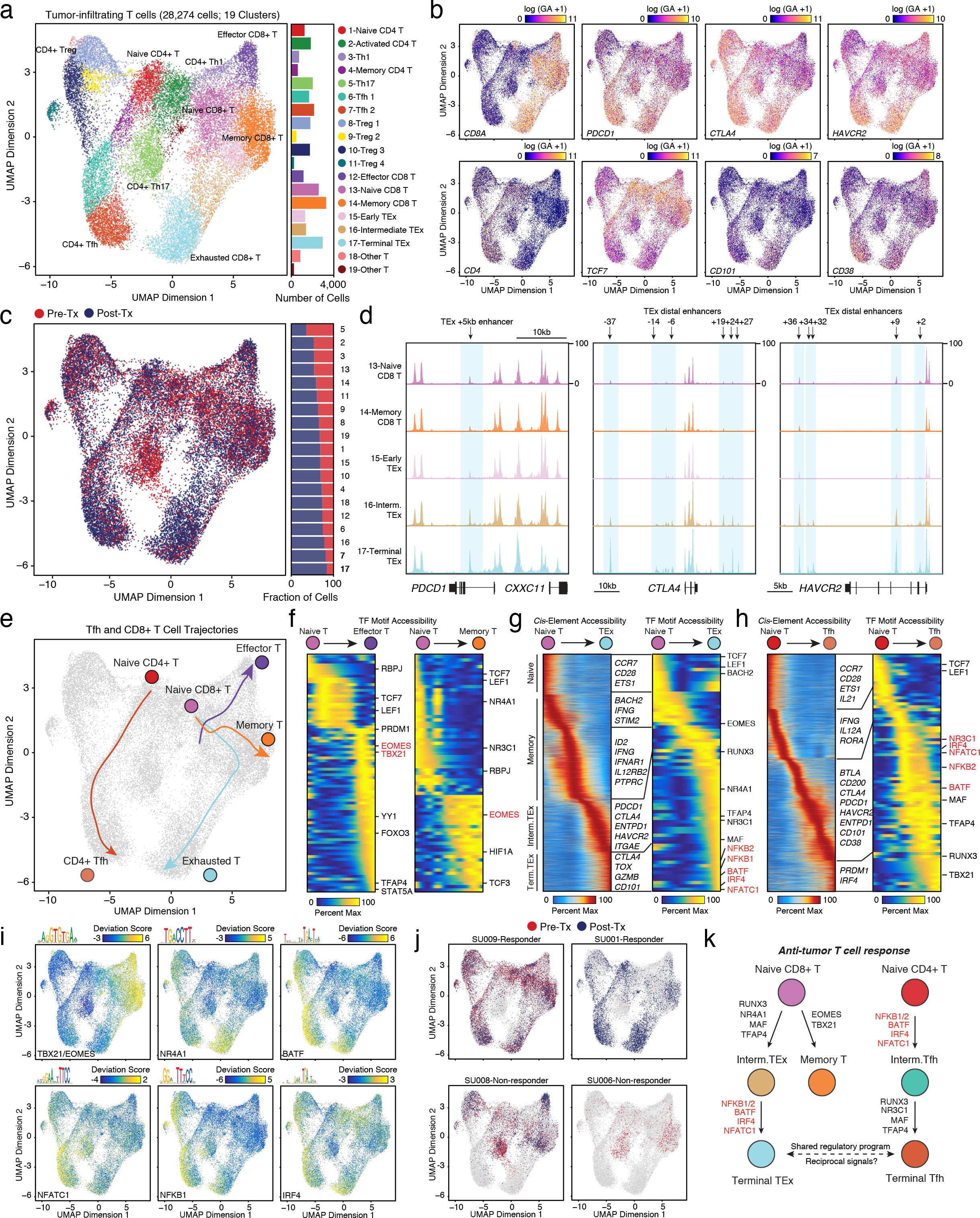
Epigenomic regulators of T cell exhaustion after PD-1 blockade. (**a**) Sub-clustering UMAP projection of 28,274 tumor-infiltrating T cells (Clusters 1-9 from tumor micro environment). scATAC-seq profiles are colored by cluster identity, as labeled on the right. Bar plot indicates the number of cells in each cluster of cells. (**b**) UMAP projection colored by log normalized gene activity scores demonstrating the accessibility of *cis*-regulatory elements linked (using Cicero) to the indicated gene. (**c**) UMAP projection of tumor-infiltrating T cells colored by pre- and post-PD-1 blockade samples. Genome tracks of aggregate scATAC-seq data, clustered as indicated in (a). Arrows indicate the position and distance (in kb) of intragenic or distal enhancers in each gene locus. (**e**) Lineage trajectories of Tfh and CD8^+^ T cell states. Lines represent double-spline fitted trajectories across pseudotime. (**f**) Pseudotime heatmap ordering of chromVAR TF motif bias-corrected deviations in effector and memory CD8^+^ T lineage trajectory. TF motifs are filtered for genes that are highly active (defined as the average percentile between total gene activity and variability > 0.75) that also demonstrate similarly dynamic gene scores across differentiation (R > 0.35 and FDR < 0.001). Heatmap of TF gene scores is shown on the right. (**g**) Pseudotime heatmap ordering of *cis*-regulatory elements (left) and chromVAR TF motif bias-corrected deviations (right) in the CD8^+^ TEx lineage trajectory. (**h**) Pseudotime heatmap ordering of *cis*-regulatory elements (left) and chromVAR TF motif bias-corrected deviations (right) in the CD4^+^ Tfh lineage trajectory. (**i**) UMAP projection of scATAC-seq profiles colored by chromVAR TF motif bias-corrected deviations for the indicated factors (**j**) UMAP projection of tumor-infiltrating T cells colored by pre- and post-PD-1 in representative individual responder and non-responsder patients. (**k**) Schematic of regulatory modules controlling TEx and Tfh differentiation.

We asked whether we could similarly group *cis*-elements according to the presence of disease-associated genetic variants, rather than TF motifs, in order to nominate cellular determinants of autoimmune disease. To achieve this, we used a list of fine-mapped causal variants associated with 21 autoimmune diseases and 18 non-immune diseases^22^ and generated a feature set for each disease that consisted of variant-containing ATAC-seq peaks. To increase the statistical power of this analysis, we also included all co-accessible elements to those containing causal variants (identified using Cicero; **Supplementary Fig. 5c**). We then measured chromatin accessibility change in all variant-associated regions genome-wide in each single cell to nominate potentially causal cell types in each disease (chromVAR; **Supplementary Fig. 5c-d**). Strikingly, this analysis revealed several distinct patterns of cell type-specific accessibility change among autoimmune diseases. As previously observed, several diseases, such as celiac disease, Type 1 diabetes, Crohn’s disease, and juvenile arthritis, primarily showed high accessibility of variant-containing putative enhancers (henceforth termed ‘variant-enhancers’) in T cell populations (**Supplementary Fig. 5d**)^22^. Other diseases, such as Kawasaki disease, multiple sclerosis, and systemic lupus erythematosus, showed high accessibility of variant-enhancers in B cells – either specifically, or in addition to accessibility in T cells – as previously described (**Supplementary Fig. 5d**)^22^. However, the comprehensive scale of scATAC-seq cell types enabled the discovery of novel patterns. For example, variant-enhancers associated with systemic sclerosis showed high accessibility in mature NK cells and pDCs, and variant-enhancers associated with ulcerative colitis showed high accessibility in mDCs and monocytes, consistent with the suggested impact of these cell types in murine models of each disease^28,29^. Additional diseases with high variant-enhancer signals in myeloid cells included a number of metabolic traits and diseases, such as fasting glucose and HDL cholesterol levels, and Type 2 diabetes, suggesting that cell-specific enhancers nominate regulatory roles for immune cells in these processes as well. The association of individual disease variants with cell type-specific enhancers could be confirmed by H3K27ac HiCHIP measurements in primary cells (**Supplementary Fig. 5e**). Altogether, these results demonstrate that scATAC-seq data may be analyzed without guidance from bulk data to identify cell types and their associated chromatin accessibility landscapes, and to examine the enrichment of these landscapes for polymorphisms associated with human disease.

### Regulatory trajectories of immune lineages

Since this dataset included progenitor and effector cell types, we asked whether the density of scATAC-seq data could be used to reconstruct cellular developmental trajectories in an unbiased manner, without the use of pre-defined markers. As a test case, we aimed to reconstruct the lineage trajectory of plasma B cell differentiation, since: (1) the entirety of this developmental program occurs in the bone marrow and blood and thus ought to be captured in our dataset, and (2) the regulatory mechanisms of this process are relatively well-defined for comparison (Fig. 3a). To achieve this, we used a nearest-neighbor approach on existing cluster definitions (Fig. 3a). We started with the plasma B cell cluster (Cluster 16) and attempted to return to the HSC cluster (Cluster 1) by sequentially selecting precursor-cell relationships with the most epigenetic similarity to the cluster of interest (computed from Euclidean distances of ATAC-seq profiles; **Methods**). For example, the nearest neighbor to the memory B cell cluster (Cluster 15) was the naïve B cell cluster (Cluster 14). The nearest neighbor to naïve B cells was the Pre-B cell cluster, and so on (Fig. 3b). Indeed, this reverse reconstruction process correctly identified the well-established cellular trajectory of plasma B cell development as the most significant trajectory among all tested trajectories (*p*<0.0002; 5,000 permutations). Finally, we generated an ordering of single cells (henceforth referred to as ‘pseudotime’) along this trajectory by computing a pseudotime vector across lineage clusters and aligning each cell to the vector in the UMAP projection (Fig. 3c **and Methods**).

We next utilized this single-cell pseudotime to identify stage-specific activities of *cis*-elements and *trans*-factors during plasma B cell differentiation. An analysis of ~10,000 *cis*-elements with dynamic accessibility patterns across the trajectory revealed specific dynamic regulatory elements near several known regulators of every stage of B cell development (Fig. 3d). For example, *cis*-elements that were accessible early in the trajectory included enhancers for *EBF1*, *RUNX1*, *IL7R*, *RAG2*, and *MEF2C*, factors that are critical for B cell lineage specification and diversity (Fig. 3d)^26,30,31^. Conversely, ~3,500 *cis*-elements that were accessible late in the trajectory included elements proximal to *PRDM1*, a critical transcription factor for plasma cell fate, and the plasma cell-specific marker *SDC1* (CD138). We also determined whether examination of chromVAR deviation scores across this developmental ordering of *trans*-factors could identify critical TFs involved in B cell development. Since chromVAR TF deviation scores calculated from scATAC-seq data can reflect the potential activity of many TFs with similar DNA-binding motifs, we integrated chromVAR deviations with dynamic *cis*-element gene scores to prune the data for the activity of specific TFs within a motif family (Fig. 3e). Indeed, this method accurately identified several previously-identified TFs that are critical for B cell differentiation (Fig. 3e). More importantly, this method could resolve fine differences in the timing of TF activity. For example, MEF2C activity was observed early in B cell development – at the stage of common lymphoid progenitors (CLPs) – consistent with its demonstrated role in lymphoid versus myeloid fate specification^31^. Immediately after the induction of MEF2C activity, we observed the sequential activity of EBF1, PAX5, and IRF4, recapitulating the known order of their physiological functions in pro-B cells, pre-B cells, and naïve B cells, respectively (Fig. 3f)^26^. Altogether, these results indicate that pseudotime ordering of scATAC-seq data can be used to accurately identify *cis*- and *trans*-regulatory factors controlling cellular differentiation.

We next applied trajectory analysis to the early stages of hematopoiesis to identify TF regulators of myeloid fate decisions, particularly of dendritic cells (DCs) – a relatively rare population of cells sparsely sampled in prior epigenomic studies. We first re-clustered 16,015 progenitor and DC scATAC-seq profiles (defined in Fig. 2) and 2,074 profiles of surface marker-defined progenitors, generated in a previous study (Fig. 3g)^11^. We identified 16 sub-clusters, and the projection of the sorted scATAC-seq profiles onto *de novo*-defined clusters revealed regulatory relationships between progenitors and significant heterogeneity in marker-defined states (Fig. 3h-i). Globally, immune lineages appeared to diverge early via three distinct branches to: (1) megakaryocyte/erythroid (Meg/E) and basophil/eosinophil (Bas/Eo) fates, (2) lymphoid fates, or (3) neutrophil/monocyte/DC fates. However, sorted progenitors did not always occupy a single *de novo*-defined regulatory state. For example, classically-defined CMPs (Lineage^−^CD34^+^CD38^+^CD10^−^CD45RA^−^CD123^mid^) were present in four *de novo*-defined clusters, including in committed pathways leading to either neutrophil/monocyte/DC fates (Clusters 2 and 11), Meg/E fates (Clusters 4-5), or Baso/Eo fates (Clusters 3-4; Fig. 3h-i). This result suggests that CMPs are heterogeneous, and that while CMPs as a population contain lineage potential for all myeloid fates, the vast majority of single cells within this population are already biased toward specific lineages. Similarly, granulocyte-macrophage progenitors (GMPs; Lineage^−^CD34^+^CD38^+^CD10^−^CD45RA^+^CD123^mid^) demonstrated a similar phenomenon and were present in four clusters downstream of the CMP (Clusters 11-14), including those leading to neutrophil differentiation, as well as previously unrecognized clusters leading to mDC and pDC fates (Fig. 3h-i).

We used pseudotime ordering to identify TF trajectories in progenitor clusters, again with a specific focus on DC development (Fig. 3j). This analysis revealed both shared and unique TF programs across myeloid fates. For example, while Meg/E and Bas/Eo progenitors shared accessibility at GATA2 motifs, Bas/Eo commitment was characterized by SPI1 (PU.1) and CEBPA motif activity, while Meg/E commitment was characterized by MYB, GATA1, and KLF1 motif activity, as previously described (Fig. 3k)^32,33^. Similarly, while neutrophil progenitors shared increased accessibility at SPI1 motifs with Bas/Eo progenitors, neutrophil commitment was accompanied by additional activity of AP-1 and CEBP motifs, and of RARA (Fig. 3k). Finally, the analysis of trajectories towards DC fates revealed three distinct possible developmental trajectories. The mDC pathway transitioned through CMP and GMP clusters, and then to Cluster 13 (monocyte-dendritic cell progenitor; MDP) and Cluster 14 (common dendritic progenitor; CDP), prior to terminal mDC differentiation. This trajectory showed accessibility at IRF8, IRF4, BCL11A, SPI1, KLF4, AP-1, and RBPJ motifs, consistent with critical roles of each of these factors in DC differentiation^34^. Importantly, there was a clear ordering of each factor’s activity in early versus late differentiation; IRF8, BCL11A, and SPI1 motifs exhibited accessibility early in CDPs, while AP-1 and RBPJ factors increased in accessibility late in terminal differentiation (Fig. 3k). For pDCs, two possible trajectories could be observed, supporting previous reports that this lineage can arise from both myeloid- and lymphoid-committed progenitors^35–37^. A first pDC trajectory transitioned directly from lymphoid-primed multipotent progenitors to differentiated pDCs, while a second trajectory traversed CMP, GMP, MDP, and CDP stages prior to pDC differentiation (Fig. 3k). Analysis of TF deviations revealed that each pathway relied on the same regulatory program, which included RUNX, IRF8, SPIB, BCL11A, and TCF4 factors, as previously demonstrated in murine and human pDCs^34^. Importantly, we did not observe significant epigenomic heterogeneity within terminal pDCs, suggesting that divergent cellular trajectories can achieve nearly identical cell states through activation of a common regulatory program.

### Single-cell chromatin landscape of intratumoral immunity

We next applied this method to primary solid tumor biopsies from BCC patients receiving PD-1 blockade. BCC is the most common cancer in humans worldwide, and recent studies demonstrated that a subset of patients with advanced BCC show significant clinical benefit from immunotherapies based on blocking the T cell inhibitory receptor PD-1^38^. However, as in many other cancers, PD-1 blockade is clinically ineffective in more than half of BCC patients^39,40^. Thus, our goal was to identify cell types that were responsive to therapy and the regulatory mechanisms controlling their activity in responder versus non-responder patients. In addition, these experiments demonstrated the feasibility of applying this method to sparse samples (as low as 500 sorted cells) from clinical biopsies.

We performed scATAC-seq on serial tumor biopsies pre- and post-PD-1 blockade (pembrolizumab) from 5 patients, plus post-therapy biopsies from 2 additional patients (total of 7 patients; 14 timepoints sampled; Fig. 4a **and Supplementary Table 3**). All patients had histologically-verified locally advanced or metastatic BCC and were poor candidates for surgical resection. To minimize non-therapy-related immune cell variation, we excluded patients with prior exposures to checkpoint blockade, or to systemic immune suppressants within 4 weeks of biopsy. We dissociated tumors into single-cell suspensions using enzymatic dissociation and sampled cells both in an unbiased fashion, analyzing total cells, or after cell sorting to enrich for T cells (CD45^+^CD3^+^), non-T immune cells (CD45^+^CD3^−^) and/or stromal and tumor cells (CD45^−^; **Supplementary Fig. 6a and Methods**). To enable pre- and post-therapy cell comparisons, biopsies were site-matched in each patient, and when possible, cells were sampled in the same manner at each timepoint. In total, we generated high-quality scATAC-seq profiles from 37,818 cells. Cells passing filter yielded on average 15 × 10^3^ unique fragments mapping to the nuclear genome, and approximately 62.5% of Tn5 insertions were within peaks present in the aggregate ATAC-seq profile from all cells, demonstrating that we could obtain high-quality scATAC-seq profiles from solid tumor biopsies (Fig. 4b **and Supplementary Fig. 6b-d**).

We clustered scATAC-seq profiles with LSI followed by SNN clustering and visualized clusters with UMAP, and identified 20 scATAC-seq clusters (Fig. 4b-c). Classification of clusters using *cis*-element accessibility and gene scores revealed a diverse ecosystem of cell types in the BCC TME, including 9 T cell clusters (high accessibility of *CD3E*, *CD8A*, and *CD4* enhancers), 2 natural killer (NK) cell clusters (high accessibility of *KLRC1* and *NCR1* enhancers), B cells and plasma cells (high accessibility of *CD19* and *SDC1* enhancers, respectively), myeloid cells that comprised mDCs and tissue macrophages (high accessibility of *CD86*, *CSF1R*, and *FLT3* enhancers), stromal endothelial cells and fibroblasts (high accessibility of *CD31* and *COL1A2* enhancers, respectively), and 4 tumor cell clusters (high accessibility of *KRT14* enhancers; Fig. 4b-d **and Supplementary Fig. 6e**). Notably, stromal and immune cells from different patients clustered together, demonstrating that these clusters did not represent patient-specific cell states or batch effects. In contrast, tumor cell clusters were largely patient-specific, consistent with prior single-cell RNA-seq studies in melanoma and head and neck cancer^41,42^, and perhaps reflecting cell state changes driven by patient-specific genome alterations (Fig. 4c). To identify potential genome alterations driving each tumor cell state, and to further support the distinction of malignant and non-malignant cells, we estimated copy number variation (CNV) from scATAC-seq data (Fig. 4e and Methods). This analysis revealed CNVs in Clusters 17-20, compared to other stromal cell populations. For example, tumor cells in patient SU010 showed ATAC-seq signal consistent with amplifications of regions of chromosomes 3 and 6, which were present in both pre- and post-therapy samples (Fig. 4e). Finally, we analyzed TF activity in each cell type and found distinct patterns of activity in immune cells, compared to stromal or tumor cells (Fig. 4f). Immune cells displayed high accessibility of TBX21, EOMES, RUNX, BCL11A, SPI1, and IRF motifs, while stromal and tumor cells displayed high accessibility of NFκB, CEBP, p63, and AP-1 motifs (Fig. 4f **and Supplementary Fig. 7a-b**). Moreover, tumor cells showed high accessibility of GLI1 motifs, consistent with the critical role of the Hedgehog signaling pathway in BCC (**Supplementary Fig. 7b**)^43^.

### Epigenetic landscape of T cell exhaustion after PD-1 blockade

We asked whether TME chromatin landscapes could identify epigenetic regulators of the anti-tumor T cell response. Since T cells can be activated by targeting either T cell inhibitory receptors (such as PD-1) or by targeting inhibitory receptor ligands (such as PD-L1) expressed on stromal cells, we examined this question in the context of both T and stromal cell populations. First, we analyzed *cis*-elements near genes encoding the known inhibitory ligands, CD47, TGFβ, and PD-L1 (Fig. 4g)^44–46^. Analysis of scATAC-seq clusters identified distinct *cis*-element patterns for each gene across stromal and tumor clusters. For example, we identified 3 differentially-accessible *cis*-elements (−35kb, +97kb, and +103kb) in the *CD47* locus, consistent with previously identified functional enhancers controlling *CD47* expression (Fig. 4g)^47^. Importantly, the tumor necrosis factor- and NFκB-responsive +97kb and +103kb enhancers were specifically accessible in tumor cells, and not stromal cells, supporting previous reports that CD47 expression on tumor cells is responsive to inflammatory signals and may mediate escape from immune surveillance^47^. Similarly, in the *TGFB1* locus, we identified 3 *cis*-elements (+29kb, +30kb, and +32kb) that were primarily accessible in stromal cells (endothelial cells and fibroblasts), consistent with the expression pattern of this gene in primary tumors (Fig. 4g)^45^. We also identified 3 *cis*-elements in the *PDL1* locus (+5kb, +9kb, and +43kb), which were previously shown to be active in 23 human cancers types in The Cancer Genome Atlas^48^. scATAC-seq data demonstrated the accessibility of these sites in tumors cells, stromal cells, and myeloid and B cells, supporting the broad expression pattern of this ligand, and suggesting that its expression may rely on the identical *cis*-regulatory elements in each cell type (Fig. 4g).

We next examined regulatory landscapes of intratumoral T cells. We first re-clustered 23,274 T cells (defined in Fig. 4) and identified 19 sub-clusters of intratumoral T cell states, revealing a rich diversity of T cell phenotypes in the TME (Fig. 5a). Classification of clusters using *cis*-element accessibility and gene scores for *CD8A* and *CD4* showed that 6 clusters represented CD8^+^ T cell states and 13 clusters represented CD4^+^ T cell states (Fig. 5b). CD8^+^ T cell states included naïve T cells (high accessibility of *CCR7 and TCF7* enhancers), effector T cells (high accessibility of *EOMES* and *IFNG* enhancers), memory T cells (high accessibility of *EOMES* enhancers, but low accessibility of effector gene enhancers), and exhausted T cells (TEx; high accessibility of *cis*-elements proximal to inhibitory receptor genes *PDCD1*, *CTLA4*, and *HAVCR2,* and to T cell dysfunction genes *CD101* and *CD38*^49^; Fig. 5b **and Supplementary Fig. 8a-b**). We also identified a cluster consistent with an intermediate TEx cluster (Cluster 16), that exhibited gene score patterns of both TEx and memory T cells (Fig. 5b). CD4^+^ T cell states included naïve T cells and several T helper cell phenotypes, such as Tregs (high accessibility of *FOXP3* and *CTLA4* enhancers), T helper 1 (Th1) cells (high accessibility of *IFNG* and *TBX21* enhancers), T helper 17 (Th17) cells (high accessibility of *IL17A* and *CTSH* enhancers), and T follicular helper (Tfh) cells (high accessibility of *CXCR5*, *IL21*, and *BTLA* enhancers; Fig. 5b **and Supplementary Fig. 8a-b**).

We focused on CD8^+^ TEx cells because previous studies demonstrated that this population is enriched for clonally-expanded tumor-specific T cells^41,50^, and that irreversibility of the TEx epigenetic state may limit the re-invigoration of tumor-specific T cells after PD-1 blockade^51^. Indeed, a comparison of pre- and post-PD-1 blockade profiles in our dataset showed that TEx cells were the most highly-expanded T cell cluster (Cluster 17) after therapy (Fig. 5c). More than 90% of TEx cells were derived from post-therapy biopsies, whereas memory and effector CD8^+^ clusters (Clusters 12 and 14) were equally derived from both timepoints. Notably, compared to other CD4^+^ T cell types, we observed an expansion of CD4^+^ Tfh cells post-therapy – nearly to the same extent as TEx cells – compared to other CD4^+^ T cell types, suggesting that PD-1 blockade impacts both CD4^+^ and CD8^+^ cell states in the TME (Fig. 5c). We first analyzed *cis*-regulatory landscapes in TEx cells to: (1) measure global changes in chromatin accessibility in TEx cells, and (2) nominate individual *cis*-elements that may regulate TEx-specific inhibitory receptor expression. Across all T cell states, we identified 35,147 *cis*-elements that were specifically accessible only in a single cluster (mean: 3,361 peaks per cluster, range: 979 peaks (naïve CD8^+^ T)-10,808 peaks (Tregs), FDR < 0.01; **Supplementary Fig. 8d**). In TEx cells, we identified 4,598 such elements, demonstrating that human T cell exhaustion is accompanied by substantial global remodeling of the chromatin accessibility landscape, and that the epigenetic state occupied by this cell state is as distinct as most other recognized T cell states, consistent with prior studies in mice^49,51,52^. Analysis of individual TEx-specific enhancers identified novel regulatory elements in inhibitory receptor loci (Fig. 5d). For example, the *PDCD1* locus (encoding PD-1) contained an intragenic *cis*-element (+5kb) with specific accessibility in TEx cells, suggesting that the persistent expression of PD-1 in exhausted human T cells is controlled by a single state-specific enhancer, and that the regulation of persistent PD-1 expression may be different in humans and mice^53^. In contrast, the *CTLA4* and *HAVCR2* loci showed TEx-specific activity of several distal *cis*-elements, compared to other CD8^+^ T cell states (Fig. 5d).

We used pseudotime ordering to identify TF trajectories in TEx differentiation, compared to effector or memory CD8^+^ T cells (Fig. 5e). Since we also observed an expansion of CD4^+^ Tfh cells post-therapy, we included this cell type in our comparison. As expected, the differentiation of naïve CD8^+^ T cells to either effector or memory cells identified the critical roles of EOMES and TBX21 (T-bet) motifs in effector and memory cell formation, once again supporting the validity of this analysis^54–56^ (Fig. 5f). Effector cell pseudotime also demonstrated the late accessibility of other known regulators, including TFAP4 and YY1, among others^57,58^. Similarly, memory cell pseudotime also showed accessibility at HIF1A and E protein sites^59^. In contrast, TEx cells showed a distinct regulatory program, which progressed through two stages of differentiation (Fig. 5g). The first stage (intermediate TEx) showed new accessibility of *cis*-elements near inhibitory receptors, as well as elements near genes associated with tissue residency, such as *ITGAE* (CD103)^60^. Accordingly, this stage of differentiation was accompanied by accessibility of NR3C1 and NR4A1 motifs, factors immediately downstream of T cell receptor (TCR) signaling^61^, as well as the RUNX3 motif, a factor that programs tissue residency of CD8^+^ T cells in tumors (Fig. 5g)^62^. The second stage (terminal TEx) showed new accessibility of cis-elements near genes associated with terminal T cell dysfunction, such as *CD101* and *TOX*^49,51^, as well as of new elements in stage 1 gene loci, such as *CTLA4* (Fig. 5g). Importantly, this stage was accompanied by accessibility of a core set of TF motifs, which included NFKB1 and NFKB2, BATF, IRF4, and NFATC1, factors that are all directly downstream of TCR signaling, and three of which have been demonstrated to play crucial roles in T cell exhaustion in mice^63–65^.

Finally, we examined the epigenetic relationship between TEx and Tfh cells. Tfh cells have previously been observed in tumors and have been proposed as a prognostic indicator of patient survival and response to checkpoint blockade^66–68^.The inferred differentiation trajectory from CD4^+^ naïve T cells to Tfh cells showed new accessibility of *cis*-elements neighboring Tfh-specific genes, such as *IL21* and *BTLA*, but also of elements near genes typically associated with TEx cells, such as inhibitory receptors, consistent with the known, but unexplained, expression of these genes in human Tfh cells (Fig. 5h **and Supplementary Fig. 8c-e**)^69^. Strikingly, differentiation was accompanied by the accessibility of Tfh regulators, but also of the same core set of TF motifs associated with TEx differentiation, including NFKB2, BATF, IRF4, and NFATC1, suggesting a common program driving the development of TEx and Tfh cells downstream of PD-1 blockade (Fig. 5h-i **and Supplementary Fig. 8f**). Indeed, the abundance of TEx and Tfh cells was similar in all patients post-therapy, and in our small cohort, was associated with therapy response (Fig. 5j). Altogether, these results map the epigenetic landscape of intratumoral TEx cells in humans and suggest that chronic TCR signals drive a shared regulatory program in TEx and Tfh cells after PD-1 blockade (Fig. 5k).

## Discussion

Recent advances in high-throughput single-cell profiling have driven new insights into cell types, RNA expression, and protein markers underlying biological processes. However, to date, single-cell chromatin accessibility profiling has been constrained by trade-offs between data quality, throughput in cell numbers, and cost per cell. Here we report a droplet-based solution for highly multiplexed single-cell chromatin accessibility profiling. The ATAC-seq profiles generated using this method are high-quality and enable an unbiased deconvolution of *cis*- and *trans*-regulatory elements underlying chromatin states with single-cell resolution. This massive scale of cell type and cell state reconstruction affords three key advantages: (1) comprehensive deconvolution of all cells in a tissue, including rare cell types and states, (2) unbiased reconstruction of cellular developmental trajectories, without the use of pre-defined markers, and (3) analysis of active regulatory DNA down to the level of individual genes and elements in single cells. The droplet-based scATAC-seq library cost is ~$0.4 per cell, has a lower multiplet rate compared to prior methods, and does not require cell sorting or non-commercial reagents.

Chromatin accessibility states of regulatory DNA encode and presage cell fates^8,70^. scATAC-seq is thus well-suited to track trajectories of cell differentiation, which can occur either as discrete steps or as a continuum of similar states. We developed a bottom-up, data-driven approach to iteratively group single cells together based on their accessible genomes, used groups of highly similar cells to reconstruct detailed cell type-specific *cis*- and *trans* regulatory maps, and highlighted disease-associated enhancers that are uniquely active in specific cell types. Moreover, the dense single-cell clusters enabled unbiased computational inference of sequential cell state transitions that could reconstruct the developmental trajectories of individual cell types, for example recapitulating decades of research on B cell and DC development. Importantly, scATAC-seq of tumor-infiltrating lymphocytes from patient biopsies identified new regulatory programs controlling T cell exhaustion and a previously unrecognized shared program with T follicular helper cells. It is tempting to speculate that this shared program may reflect an evolutionarily conserved pathway to synchronize CD4^+^ and CD8^+^ T cell responses to chronic pathogen infection, such that CD4^+^ Tfh cells support antibody formation as well as long-term activation of CD8^+^ T cells, perhaps through IL-21^71–73^. Nevertheless, future studies targeting these regulatory pathways may nominate therapeutic interventions that synergize with PD-1 blockade in cancer. In summary, we describe a method for generating large-scale single-cell chromatin accessibility profiles on a widely-distributed single-cell investigation platform, enabling unbiased discovery of cell types and regulatory DNA elements in complex tissues.

## Acknowledgments

We thank members of the Chang and Greenleaf laboratories and 10x Genomics for helpful discussions. We thank the following people at 10x Genomics: Alaina Puleo for sorting cells, John Chevillet for training, Zachary Bent and Michael Dodge for reagents development, Rachel Gerver and Wei Wang for microfluidics, and Abby Gallegos, Alvaro Gonzales, Nikka Keivanfar, Shamoni Maheshwari, Patrick Marks, Jeff Mellen, Rudy Rico, and Kevin Wu for computational and software support. We thank Xuhuai Ji, Dhananjay Wagh, and John Coller at the Stanford Functional Genomics Facility, and Charles Bruce at 10x Genomics for sequencing support and Alexander Valencia for assistance with clinical specimen processing. This work was supported by the National Institutes of Health (NIH) grants P50HG007735 (H.Y.C. and W.J.G.), K08CA23188-01 (A.T.S.), K99-AG059918 (M.R.C.), and S10OD018220 (Stanford Functional Genomics Facility), the Parker Institute for Cancer Immunotherapy (A.T.S. and H.Y.C.), the Michelson Foundation (A.T.S.), and the Scleroderma Research Foundation (H.Y.C.). A.T.S. was supported by a Bridge Scholar Award from the Parker Institute for Cancer Immunotherapy and a Career Award for Medical Scientists from the Burroughs Wellcome Fund. K.E.Y. was supported by the National Science Foundation Graduate Research Fellowship Program (NSF DGE-1656518) and a Stanford Graduate Fellowship. W.J.G is a Chan Zuckerberg Biohub investigator. H.Y.C. is an investigator of the Howard Hughes Medical Institute.

## Author Contributions

A.T.S., J.M.G., G.X.Y.Z., W.J.G., and H.Y.C. conceived the project. A.T.S, J.M.G., Y.Q., K.E.Y., M.R.C., M.R.M, S.E.P., F.M., G.P.M. J.C.B., D.J., C.M.N., J.W., Y.Y. performed experiments. J.M.G. lead the analysis of scATAC-seq data. B.N.O., P.S. and L.W. contributed to the Cell Ranger ATAC software and contributed to data analysis with P.G.G. A.L.S.C. obtained clinical specimens. A.T.S., H.Y.C, W.J.G., and G.X.Y.Z. guided experiments and data analysis. A.T.S, J.M.G., G.X.Y.Z., W.J.G., and H.Y.C. wrote the manuscript with input from all authors.

## Conflicts of Interest

H.Y.C. is a co-founder of Accent Therapeutics and Epinomics and is an adviser to 10x Genomics and Spring Discovery. W.J.G is a co-founder of Epinomics and an adviser to 10x Genomics, Guardant Health, and Centrillion. F.M., G.P.M., B.N.O., P.S., J.C.B., D.J., C.M.N., J.W., L.W., Y.Y., P.G.G., and G.Y.Z. are employees of 10x Genomics. A. L. S. C. was an advisory board member and clinical investigator for studies sponsored by Merck, Regeneron, Novartis, Galderma, Genentech Roche. Stanford University holds patents on ATAC-seq, on which P.G., W.J.G., and H.Y.C. are named as inventors.

## Methods

### Human subjects

This study was approved by the Stanford University Administrative Panels on Human Subjects in Medical Research, and written informed consent was obtained from all participants.

### Cell lines and PBMC/BM samples

Human (GM12878) and Mouse A20 (ATCC TIB-208) B Lymphocytes were acquired and cultured according to guidelines from Coriell and ATCC, respectively. Fresh PBMCs, GM12878, and A20 cells were frozen according to the instructions outlined here: https://assets.ctfassets.net/an68im79xiti/2ptJYphPcPGfSPisq0cVuu/c8a83f93383c2fd1ce7cc49abc837992/CG000169_DemonstratedProtocol_NucleiIsolation_ATAC_Sequencing_Rev_B.pdf. Briefly, PBMCs were cryopreserved in IMDM + 40% FBS + 15% DMSO. GM12878 and A20 cells were cryopreserved in RPMI + 15% FBS + 5% DMSO. For the monocyte and T cell mixing experiments, nuclei were first extracted and transposed, then mixed at indicated ratios. To avoid pipetting errors, a large number of nuclei were mixed after nuclei extraction and transposition, and a smaller number of nuclei were loaded onto the microfluidics chip for scATAC library generation. We also performed a similar mixing experiment using naïve and memory T cells (**Supplementary Table 1**), which performed similarly and is included in the available data section.

Healthy volunteer PBMC and BM samples were obtained from AllCells or the Stanford Blood Center (SBC). Mononuclear cells from each sample were isolated by Ficoll separation, resuspended in 90% FBS + 10% DMSO, and cryopreserved in IMDM + 40% FBS + 15% DMSO. Samples were then thawed at 37°C for 5 min and resuspended in media prior to cell enrichment using magenetic-activated cell sorting (MACS) or FACS (**Supplementary Table 2**). All MACS-enriched populations were obtained from AllCells and isolated per manufacturer recommendations (as outlined in **Supplementary Table 2**). FACS-isolated populations were obtained from AllCells or SBC and sorted as follows. CD4^+^ T helper cells were sorted as naive T cells (CD4^+^CD25^−^CD45RA^+^) or memory T cells (CD4^+^CD25^−^CD45RA^−^) using the following antibodies: anti-human CD45RA-PERCPCy5.5 (Clone HI100, Biolegend), anti-human CD4-APC-Cy7 (Clone OKT4, Biolegend), and anti-human CD25-FITC (Clone BC96, Biolegend). Dendritic cells and basophils were sorted as CD3^−^CD19^−^CD11c^+^HLA-DR^+^ (DCs) and CD3^−^CD19^−^CD123^+^ (basophils) using the following antibodies: CD11C-PECy7 (Clone B-ly6, BD), HLA-DR-APCCy7 (Clone G46-6, BD), CD123-BV421 (Clone 6H6, Biolegend), CD3-FITC (Clone UCHT1, BD), and CD19-AlexaFluor 488 (Clone HIB19, Biolegend). All antibodies were validated by the manufacturer in human peripheral blood samples, used at a 1:200 dilution, and compared to isotype and no staining control samples.

### BCC sample collection and cell sorting

Fresh BCC biopsies were digested in 5 mL DMEM/F12 media + 250 μg/mL Liberase TL and 200 U/mL DNAse I with the gentleMACS Octo system at 37°C for 3 hours at 20 rpm. After tissue pieces were fully digested, 50 μL of 500 mM EDTA was added and samples were collected by centrifugation at 300xg for 5 minutes. Single-cell suspensions were filtered through 70 μm mesh and pelleted by centrifugation at 300xg at 4°C for 10 minutes. Finally, cells were then resuspended in 1 mL of RPMI media and cryopreserved in FBS supplemented with 10% DMSO.

Cells were gently thawed at 37°C for 5 min and resuspended in media prior to FACS. Cells were stained with anti-CD45 V500 (clone HI30, cat. no. 560779, BD Biosciences), anti-CD3 FITC (clone OKT3, cat. no. 11-0037-41, Invitrogen), anti-CD8 Pacific Blue (clone 3B5, cat. no. MHCD0828, Invitrogen), anti-PD-1 APC/Cy7 (clone EH12.2H7, cat. no. 329921, BioLegend), and anti-HLA-DR eVolve 605 (clone LN3, cat. no. 83-9956-41, Affymetrix-Ebioscience). All antibodies were used at a 1:200 dilution, with the exception of anti-CD45 and anti-HLA-DR antibodies, which were used at a 1:100 dilution. Propidium iodine (cat. no. P3566, Invitrogen) was used for live/dead staining at a final concentration of 2.5 μg/mL. PI-negative live cells were sorted as T cells (CD45+CD3+), non-T immune cells (CD45+CD3-), or tumor/stromal cells (CD45-CD3-) and further processed using scATAC-seq.

### Single-cell ATAC-Seq using the 10x Chromium platform

All protocols to generate scATAC-seq data on the 10x Chromium platform, including sample prep, library prep, instrument and sequencing settings, are described below and are also available here: https://support.10xgenomics.com/single-cell-atac.

#### Nuclei Isolation

The isolation, washing, and counting of nuclei suspensions were performed according to the Demonstrated Protocol: Nuclei Isolation for Single Cell ATAC Sequencing (10x Genomics). Briefly, anywhere from 100,000 to 1,000,000 cells were added to a 2 ml microcentrifuge tube and centrifuged (300 rcf for 5 min at 4°C). The supernatant was removed without disrupting the cell pallet and 100 *µ*l chilled Lysis Buffer (10 mM Tris-HCl (pH 7.4); 10 mM NaCl; 3 MgCl_2_; 0.1% Tween-20; 0.1% Nonidet P40 Substitute; 0.01% Digitonin and 1% BSA) was added then pipette mixed 10 times. The microcentrifuge tube was then incubated on ice, with the length of time optimized for each cell type: GM12878 and A20 cell lines were 5 min; PB and BM cells were 3 min. Following lysis, 1 mL of chilled Wash Buffer (10 mM Tris-HCl (pH 7.4); 10 mM NaCl; 3 MgCl_2_; 0.1% Tween-20 and 1% BSA) was added and the resulting solution and pipette mixed 5 times. Nuclei were centrifuged (500 rcf for 5 min at 4°C) and the supernatant removed without disrupting the nuclei pellet. Based on the starting number of cells and desired final nuclei concentration, an appropriate volume of chilled Diluted Nuclei Buffer (10x Genomics; 2000153) was used to resuspend nuclei. The resulting nuclei concentration was determined using a Countess II FL Automated Cell Counter. Nuclei were then immediately used to generate single cell ATAC-seq libraries as described below.

#### Library Construction

scATAC-seq libraries were prepared according to the Chromium Single Cell ATAC Reagent Kits User Guide (10x Genomics; CG000168 Rev B). Briefly, the desired number of nuclei were combined with ATAC Buffer (10x Genomics; 2000122) and ATAC Enzyme (10x Genomics; 2000123/2000138), to form a Transposition Mix which was then incubated for 60 min at 37°C. A Master Mix comprising of Barcoding Reagent (10x Genomics; 2000124), Reducing Agent B (10x Genomics; 2000087) and Barcoding Enzyme (10x Genomics; 2000125/2000139) was then added to the same tube as Transposed Nuclei. The resulting solution was loaded onto a Chromium Chip E (10x Genomics; 2000121) in a Chip Holder (10x Genomics; 330019). Vortexed Chromium Single Cell ATAC Gel Beads (10x Genomics; 2000132) then Partitioning Oil (10x Genomics; 220088) were also loaded onto the same Chromium Chip E before attaching a 10x Gasket (10x Genomics; 370017/3000072) and placing into on a Chromium™ Single Cell Controller instrument (10x Genomics, Pleasanton, CA, USA). Resulting single-cell GEMs were collected at the completion of the run (~7 min) and linear amplification was performed in a C1000 Touch Thermal cycler with 96-Deep Well Reaction Module (Bio-Rad; 1851197): 72°C for 5 min, 98°C for 30 s, cycled 12 x: 98°C for 10 s, 59°C for 30 s and 72°C for 1 min. Emulsions were coalesced using tje Recovery Agent (10x Genomics; 220016) then subjected to Dynabeads (2000048) and SPRIselect reagent (Beckman Coulter; B23318) bead clean-ups. Indexed sequencing libraries were constructed by combining the barcoded linear amplification product with a Sample Index PCR Mix comprising of SI-PCR Primer B (10x Genomics; 2000128), Amp Mix (10x Genomics; 2000047/2000103) and Chromium i7 Sample Index (10x Genomics; 3000262). Amplification was performed in a C1000 Touch Thermal cycler with 96-Deep Well Reaction Module: 98°C for 45 s, cycled variable amounts depending on cell load: 98°C for 20 s, 67°C for 30 s, 72°C for 20 s with a final extension of 72°C for 1 min. The barcode sequencing libraries were subjected to a final bead clean-up SPRIselect reagent and quantified by quantitative PCR (KAPA Biosystems Library Quantification Kit for Illumina platforms; KK4824). Sequencing libraries were loaded on an Illumina sequencer with 2 × 50 paired-end kits using the following read length: 50 bp Read 1N, 8 bp i7 Index, 16 bp i5 Index and 50 bp Read 2N.

| Instrument | Loading Concentration (pM) | PhiX (%) |
| --- | --- | --- |
| NextSeq 500 | 1.7 | 1 |
| HiSeq 2500 (RR) | 11 | 1 |
| HiSeq 4000 | 180 | 1 |
| NovaSeq | 250 | 1 |

#### Oligonucleotide sequences

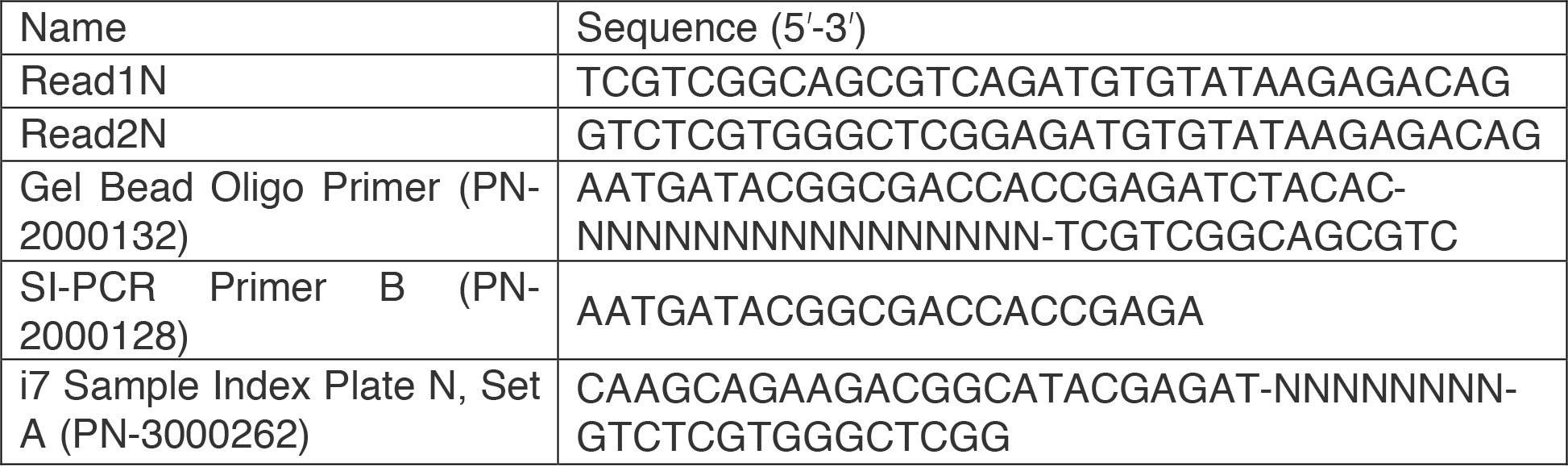

### Availability of Data Processing and Analysis Software

All data processing steps and methods used in the manuscript are described in detail below. We also have designed and made the following tools freely available:

Cell Ranger ATAC: This software performs initial data processing of scATAC sequencing reads (including de-multiplexing, genome alignment, and read de-duplication), as described below and used in this manuscript. This software will also perform additional downstream analysis, including the identification of open chromatin regions, motif annotations, and differential accessibility analysis, similar to what was performed in this manuscript and described below. https://support.10xgenomics.com/single-cell-atac/software/pipelines/latest/what-is-cell-ranger-atac

Loupe Cell Browser: This is an interactive visualization software that shows ATAC-seq peak profiles for scATAC-seq cell clusters, similar to the analysis done in this manuscript and described below. https://support.10xgenomics.com/single-cell-atac/software/visualization/latest/what-is-loupe-cell-browser

### Data Processing using Cell Ranger ATAC Software

The Cell Ranger (CR) Software (version 1.0; https://support.10xgenomics.com/single-cell-atac/software/pipelines/latest/algorithms/overview) was used for alignment, de-duplication, and identification of transposase cut sites. First, the 16bp barcode sequence was processed to fix the occasional sequencing error in barcodes. Barcode sequences were obtained from the I2 index reads. An observed barcode not present in the whitelist of barcodes can be corrected to a whitelist barcode if it is within 2 hamming distance away, and has >90% probability of being the real barcode (based on the abundance of the barcode and quality value of incorrect bases). Then, the cutadapt tool was used to identify and trim any adapter sequence in each read. Third, the trimmed read pairs were aligned to a reference using BWA-MEM with default parameters. Reads less than 25bp are not aligned and flagged as unmapped. Fragments are identified as read pairs with MAPQ>30, non-mitochondria reads and not chimerically mapped. The start and end of the fragments are adjusted (+4 for + strand and −5 for - strand) to account for the 9bp region the transposase enzyme occupies during the transposition. Lastly, fragments with identical start and end positions were counted once. The most common barcode sequence is assigned to the fragments, with ties broken by picking the barcode sequence with the highest read counts. One of the read-pairs with that barcode sequence is labeled as the ‘original’ and the other read-pairs in the group are marked as duplicates of the fragment in the BAM file.

### scATAC-seq Data Analysis

#### Filtering Cells by TSS Enrichment and Unique Fragments

Enrichment of ATAC-seq accessibility at TSS was used to quantify data quality without the need for a defined peak set. Calculating enrichment at TSS was performed as previously described^48^, and TSS positions were acquired from the Bioconductor package from “TxDb.Hsapiens.UCSC.hg19.knownGene”. Briefly, Tn5 corrected insertions were aggregated +/− 2,000 bp relative (TSS strand-corrected) to each unique TSS genome wide. Then this profile was normalized to the mean accessibility +/− 1,900-2,000 bp from the TSS and smoothed every 51bp in R. The calculated TSS enrichment represents the max of the smoothed profile at the TSS. We then filtered all single cells that had at least 1,000 unique fragments and a TSS enrichment of 8 for all data sets.

#### Generating a Counts Matrix

To make a cell by feature counts matrix, we first read each fragment into R using readr. Next, we converted these fragment GenomicRanges into Tn5 insertion GenomicRanges by concatenating GenomicRanges for each “start” and “end” of the fragments (1bp width). Next, we used “findOverlaps” to find all overlaps with the feature by insertions. Then we added a column with the unique id (integer) cell barcode to the overlaps object and fed this into a sparseMatrix in R. To calculate the fraction of Tn5 insertions in peaks, we used the colSums of the sparseMatrix and divided it by the number of insertions for each cell id barcode using “table” in R. The counts matrix was then log-normalized by using edgeR’s “cpm(matrix, log = TRUE, prior.count = 3)” in R. The prior count is used to lower the contribution of variance from elements with lower count values as previously described. This normalization assumes that global differences in accessibility are minor to relative differences by depth normalizing within accessible regions.

#### Generating Union Peak Sets with Latent Semantic Indexing

We created a union peak set by adapting a previous workflow as follows^12^. Prior to calling peaks, we constructed 2.5kb windows that were tiled across the genome by using “tile(hg19chromSizes, width = 2500)” in R. Next, a cell by window sparse matrix was computed by counting the Tn5 insertion overlaps for each cell using “findOverlaps” in R as described above. This matrix was then binarized and pruned to the top 20,000 most accessible sites across all cells. We then reduced the dimensionality as previously described by computing the term frequency-inverse document frequency (“TF-IDF”) transformation^9^. Briefly we divided each index by the colSums of the matrix to compute the cell “term frequency”. Next we multiplied these values by log(1 + ncol(matrix) / rowSums(matrix)) which represents the “inverse document frequency”. This normalization resulted in a TF-IDF matrix that was then used as input to irlba’s singular value decomposition (SVD) implementation in R. We then retained only the 2nd-25^th^ dimensions (1^st^ dimension is associated with cell read depth^12^) and created a Seurat object and identified crude clusters using Seurat’s SNN graph clustering (v2.3) with “FindClusters” with a default resolution of 0.8. If the minimum cluster size was below 200 cells, the resolution was decreased until this criterion was reached leading to a final resolution of 0.8^N^ (where N represents the iterations until the minimum cluster size is 200 cells).

Peak calling for each cluster was performed independently to get high-quality, fixed-width peaks that represent the epigenetic diversity of all samples based on previous work^48^. For each cluster, peak calling was performed on Tn5-corrected single-base insertions (each end of the Tn5-corrected fragments) using the MACS2 callpeak command with parameters “--shift -75 --extsize 150 --nomodel --call-summits --nolambda --keep-dup all -q 0.05.” The peak summits were then extended by 250bp on either side to a final width of 501bp, filtered by the ENCODE hg19 blacklist (https://www.encodeproject.org/annotations/ENCSR636HFF/), and then filtered to remove peaks that extend beyond the ends of chromosomes.

Overlapping peaks called within a single sample were handled using an iterative removal procedure as previously described^48^. First, the most significant peak is kept and any peak that directly overlaps with that significant peak is removed. Then, this process iterates to the next most significant peak and so on until all peaks have either been kept or removed due to direct overlap with a more significant peak. This was performed on each cluster’s peak set and then the top 100,000 extended summits (ranked by MACS2 score) were kept to arrive at a “cluster-specific peak set” for each cluster. We then normalized the MACS2 peak scores (−log10(q-value)) for each sample and converted them to a “score quantile” by converting each individual score to a quantile using “trunc(rank(v))/length(v)” in R (where v represents the vector of MACS2 peaks scores). This normalization allows for direct comparison of peaks across clusters, enabling the generation of a union peak set for each dataset.

We next compiled a union peak set containing the important peaks observed across all clusters. First, all cluster peak sets were combined into a cumulative peak set and trimmed for overlap using the same iterative procedure mentioned above. Again, this procedure keeps the most significant (in this case, score quantile) peak and discards any peak that overlaps directly with the most significant peak. Lastly, we removed any peaks that spanned a genomic region containing “N” nucleotides and any peaks mapping to the Y chromosome.

#### Reads-in-peaks-normalized bigwigs and sequencing tracks

To visualize ATAC-seq cluster data genome-wide, we created ATAC-seq signal tracks that have been normalized by the number of reads in peaks, as previously described^48^. Briefly, we created fragment files that contained all cells belonging to a specific cluster and then counted the number of Tn5 insertions in the corresponding peak set. The number of Tn5 insertions were computed in windows genome-wide using “slidingWindows(chromSizes,100,100)”. Next, we created a run-length encode using “coverage” in R and normalized the total reads to a scale factor that normalizes the reads-in-peaks to 10 million reads within peaks. This object was then converted into a bigwig using rtracklayer “export.bw” in R. For plotting tracks, the bigwigs were read into R using rtracklayer “import.bw(as=”Rle”)” and plotted within R or visualized with WashU Epigenome browser. All track figures in this paper show groups of tracks with matched normalized y-axis scales.

To visualize scATAC-seq data, we read the fragments into a GenomicRanges object in R. We then computed sliding windows across each region we wanted to visualize every 100 bp “slidingWindows(region,100,100)”. We computed a counts matrix for Tn5-corrected insertions as described above and then binarized this matrix. We then returned all non-zero indices from the matrix (cell × 100bp intervals) and plotted them in ggplot2 in R with “geom_tile”.

#### ATAC-seq-centric Latent Semantic Indexing clustering and visualization

We clustered scATAC-seq data using an approach that does not require bulk data or prior knowledge. To achieve this, we adopted the strategy by Cusanovich et. al^9^, to compute the term frequency-inverse document frequency (“TF-IDF”) transformation. Briefly we divided each index by the colSums of the matrix to compute the cell “term frequency.” Next, we multiplied these values by log(1 + ncol(matrix) / rowSums(matrix)), which represents the “inverse document frequency.” This resulted in a TF-IDF matrix that was used as input to irlba’s singular value decomposition (SVD) implementation in R. We then used the first 50 reduced dimensions as input into a Seurat object and then crude clusters were identified by using Seurat’s (v2.3) SNN graph clustering “FindClusters” with a default resolution of 0.8. We found that there was detectable batch effect that confounded further analyses. To attenuate this batch effect, we calculated the cluster sums from the binarized accessibility matrix and then log-normalized by using edgeR’s “cpm(matrix, log = TRUE, prior.count = 3)” in R. Next, we identified the top 25,000 varying peaks across all clusters using “rowVars” in R. This was done on the cluster log-normalized matrix vs the sparse binary matrix because: (1) it reduced biases due to cluster cell sizes, and (2) it attenuated the mean-variability relationship by converting to log space with a scaled prior count. These 25,000 variable peaks were then used to subset the sparse binarized accessibility matrix and recomputed the “TF-IDF” transform. We used singular value decomposition on the TF-IDF matrix to generate a lower dimensional representation of the data by retaining the first 50 dimensions. We then used these reduced dimensions as input into a Seurat object and then crude clusters were identified by using Seurat’s (v2.3) SNN graph clustering “FindClusters” with a default resolution of 0.8. These same reduced dimensions were used as input to Seurat’s “RunUMAP” with default parameters and plotted in ggplot2 using R.

For sub-clustering analyses (Hematopoiesis: CD34^+^ BM and DCs; Tumor: T cells), we computed the cluster sums again and log-normalized using edgeR’s “cpm(matrix, log = TRUE, prior.count = 3)” in R. We identified the top 10,000 and 5,000 varying peaks for CD34^+^ cells and T cells, respectively. These variable peaks were then used to subset the sparse binarized accessibility matrix and recompute the “TF-IDF” transform. We then used singular value decomposition on the TF-IDF matrix to generate a lower dimensional representation of the data by retaining the first 25 dimensions. We then used these reduced dimensions (1-25 and 2-25, respectively) as input into a Seurat object, and then crude clusters were identified by using Seurat’s (v2.3) SNN graph clustering “FindClusters” with a default resolution of 0.8. These same reduced dimensions were used as input to Seurat’s “RunUMAP” and plotted in ggplot using R.

#### Inferring copy number amplification

To infer DNA copy number amplifications from scATAC-seq data, we first tiled the genome into 10-Mb windows using “slidingWindows” of GenomicRanges for chromosome sizes in R with a step size of 2Mb. These window positions were then filtered against regions with known artefactual mapping issues using the ENCODE hg19 blacklist with the “setdiff” function in R. Then, a cell by window binarized matrix was constructed, as described above. Next, the insertions per bp was determined within each filtered 10-Mbp window. The percent GC content was computed for each filtered 10-Mbp window using the hg19 BSgenome in R. To estimate if a region is amplified, we identified the 100 nearest neighbors based on GC content and computed the average log2(fold change). If this was above 1 we considered this region as a candidate for amplification. By looking at the median log2(fold change) for each patient across each cluster we tried to identify amplified regions.

#### Transcription factor footprinting

We wanted to characterize relative TF occupancy through TF footprints, as previously described^48^. For each peak set, we used CIS-BP motifs (from chromVAR motifs human_pwms_v1) to calculate motif positions using motifmatchr “matchMotifs(positions = “out”)”. Next, we computed the Tn5 bias for each sample by constructing hexamer bias table using “oligonucleotidefrequency” function from Biostrings in R. Then, we calculated a hexamer table for each TF by counting the hexamers relative to each stranded motif position +/−250 bp from the motif center. Using the sample’s hexamer frequency table, we could then compute the expected Tn5 insertions by multiplying the hexamer position frequency table by the observed/expected Tn5 hexamer frequency.

To assess the reproducibility of footprints, we subsampled each cluster fragments 2 times at a sampling rate of 60% to have maximum variability. To calculate the insertions around these sites, we converted the Tn5-corrected insertions GenomicRanges (see above) into a coverage run-length encoding using “coverage”. For each individual motif, we iterated over the chromosomes, computing a “Views” object using “Views(coverage, motif positions)”. This “Views” object was converted to a matrix using “as.matrix” which was then used to calculate the following and the colSums for “-stranded” motifs were reversed and the colSums for NOT “-stranded” motifs were summed. To better compare footprints across samples, we normalized these footprints by the mean values +/−200-250 bp from the motif center. Next, we divided the footprints by the expected Tn5 bias to attempt to account for the inherent Tn5 bias. While this strategy is effective, it does not fully account for all of Tn5’s sequence bias. We then plot the mean and standard deviation for each footprint pseudo-replicate.

#### ChromVAR

In addition to TF footprinting, we measured global TF activity using chromVAR^4^. We used as input the raw insertion counts for all peaks and the CIS-BP motif (from chromVAR motifs “human_pwms_v1”) matches within these peaks from motifmatchr. We then computed the GC bias-corrected deviation scores using the chromVAR “deviationScores” function. All plots used the “deviationScores” in R and variability was computed by using “rowVars” in R.

#### Computing Gene Activity Scores using Cicero Co-Accessibility

We calculated gene activities using the R package Cicero as previously described^20^. We first used the sparse binary matrix and created a cellDataSet, detectedGenes, and estimatedSizeFactors. Next, we created a “cicero_cds” with k=50 and the “reduced_coordinates” being the corresponding UMAPs. This function returns aggregated accessibility across groupings of cells based on nearest-neighbor rules. We then identified all peak-peak linkages that were within 250 kb by resizing the peaks to 250 kb and then overlapping them with the peak summits/centers. We removed all duplicates and same peak-peak links. Next, we calculated the pearson correlation for each peak-peak link and created a connections data.frame where the first column is peaki and the second column is peakj and third coaccessibility (pearson correlation). We then created a gene data.frame by getting genes from the TxDb

TxDb “TxDb.Hsapiens.UCSC.hg19.knownGene” in R. We altered the start of “MEF2C” to 88014057 because this alternative transcript start site is where we see strong promoter accessibility. We then resized each gene to its TSS and created a window +/− 2.5 kb from the TSS and then annotated our “cicero_cds” using “annotate_cds_by_site”. We then calculated gene activities by using “build_gene_activity_matrix” with a coaccess cutoff of 0.35. Lastly we normalized the gene activities by using “normalize_gene_activities” and the read depth of the cells. We wanted to adapt these activity scores to be more interpretable as pseudo RNA-seq so we further log normalized them by computing “log2(GA*1000000 +1)” which we refer to as further in the text as log2(GA + 1) (analogous to logCPM transformation). This conversion made the scores more interpretable and allowed them to be used further in TF deduplication and cell annotations.

#### GWAS PICs Causal Autoimmune Variants using Cicero Co-Accessibility and chromVAR

We sought to characterize cell-type specific enrichments in known disease associated regulatory elements. To do this analysis, we downloaded the causal SNPs for 39 diseases from http://pubs.broadinstitute.org/pubs/finemapping/dataportal.php. We then converted these into a GenomicRanges object and overlapped them with ATAC-seq peaks. We wanted to increase our power beyond direct overlaps, thus we used our cicero co-accessibility links to extend direct overlaps to peaks that are also co-accessible. To do this analysis, for every peak that had an overlap, we took all peaks that were co-accessible above 0.35 and then marked them as overlapping. We then created an overlap matrix for every group of SNPs partitioned by disease. We then used this as input to chromVAR’s “computeDeviations” with the scATAC-seq raw insertion counts for all peaks. We then used the “deviationScores” from chromVAR and plotted the median score across each cluster in R.

#### HiChIP MetaV4C support for Cicero Co-Accessibility Links

We wanted to further support predicted Cicero co-accessibility links using previously published 3D chromosome conformation data as previously described^48^. Briefly, we used published histone H3K27ac HiChIP data from primary T cell subsets^23^ (Naive, Th17, and Treg) to support our predicted Cicero co-accessibility links. First, we converted our links to 10kb resolution by flooring each coordinate (gene start and peak center) to the nearest 10kb window and deduplicated. To make distance-scaled Meta-Virtual 4C plots, each chromosome was retrieved from the “.hic” interactions file using juicer dump at 10-Kb resolution and read into a “sparseMatrix” in R (each coordinate in the matrix corresponding to a 10-Kb interaction bin). Then, for each peak-to-gene link longer than 100kb, the upstream or downstream window (depending on the peak’s location relative to the TSS) was identified and then interpolated linearly using the “approx” function to get the value at each 0.1%. This was then summed for each predicted interaction and divided by the total number of predicted interactions. Replicate reproducibility was visualized with the mean profile shown as a line and the shading surrounding the mean representing the standard deviation between replicates. Lastly, we wanted to test the specificity of our scATAC-seq T cell clusters in Naïve, Th17 and Treg H3K27ac HiChIP. We computed the cluster sums for each cluster from the binarized accessibility matrix for clusters 21 (Naïve), 24 (Memory) and 25 (Treg), log-normalized and computed the row-wise z-scores. We then took the top 25,000 peaks by Z-score for each cluster and then overlapped these with the cicero links. We then plotted the metav4C for each of the 3 subtypes with replicate reproducibility as described above.

#### Overlap of Cicero Co-Accessiblity Links with GTEx eQTLs

eQTLs from the Genotype-Tissue Expression project were used to support the ATAC-seq defined Cicero co-accessibility links as previously described^48^. First, we identified all gene starts from gencode v19 (https://gtexportal.org/home/datasets) and extended them +/− 2.5kb, as we did when computing the gene activities, and then overlapped all peaks with these regions using findOverlaps. We then labeled peaks that overlapped with the extended gene starts as a promoter peak and identified all ATAC-seq peak-to-promoter links. We chose to do this analyses with gencode v19 because we wanted the genes to match with the eQTL data set. GTEx eQTL data (version 7) was downloaded from https://gtexportal.org/home/datasets and the *.signif_variant_gene_pairs.txt.gz files were used. All eQTLs located more than 250kb away from the predicted gene pair were removed to maintain consistency with the 250kb window used in predicting ATAC-seq peak-to-promoter links. Next, the nearest genes for each eQTL were determined using “distanceToNearest” with the eQTL regions to all gene starts in gencode v19 in R. Then, all eQTLs that were paired with the nearest gene were removed to better test the predictive power of the non-nearest gene peak-to-gene link predictions. All peak-to-gene links were then overlapped with these filtered eQTLs using “findOverlaps” and then matched based on the predicted linked gene. To assess the significance of these overlaps, we created 250 random peak-to-gene link sets by taking all peaks from the hematopoiesis union peak set and randomly assigning these peaks to any gene within 250 kb of the peak summit. Then, we calculated the z-score and enrichment of our determined peak-to-gene links compared to the randomized peak sets. We then calculated the adjusted p-value using the Benjamini-Hochberg correction.

#### Constructing ATAC-seq Pseudo-Bulk Replicates of Maximal Variance

We wanted to perform analyses that treated each cluster as a bulk ATAC-seq sample, but required a method that could create replicates that convey close to the true population variance within a cluster and potential batch effects. For each cluster, we first checked whether each cluster contained 2 or more individual 10x runs that had at least 100 cells within the cluster. If true, then individual 10x runs cells were summed from the binarized matrix (maximum of 500 cells randomly sampled) to create pseudo-bulk replicates. If this condition was not true for the cluster, and there was at least one 10x run that had at least 100 cells and the remaining cells were at least 100 cells then one 10x run was summed from the binarized matrix (maximum of 500 cells randomly sampled) to create a pseudo-bulk replicate. Then for the remaining cells a pseudo-bulk replicate was constructed by randomly sampling 100 cells from the remaining cells 250 times and we kept the sampling that produced the highest within cluster total log-variance (cpm(mat, log=TRUE, prior.count=3) then summed “rowVars” with the 1 10x run replicate). If neither of these conditions were met, then two replicates of 100 cells were randomly sampled 250 times and we kept the sampling that produced the highest within cluster total variance (cpm(mat, log=TRUE, prior.count=3) then summed “rowVars” with the both sampled replicates). Lastly, if the cluster was smaller than 150 total cells, the number of cells sampled was 2/3 the size of the cluster. This workflow was designed to construct pseudo-replicates from single-cell clusters that produced high variance to attempt to capture true biological variation. In general, this approach is still an underestimate of variation when fully simulating replicates, so it’s important to be conservative when using these pseudo-replicates in further analyses.

#### Constructing Gene Activity Pseudo-Bulk Replicates of Maximal Variance

We wanted to perform analyses that treated each cluster as a bulk ATAC-seq sample, but required a method that could create replicates that convey close to the true population variance within a cluster and potential batch effects. For each cluster, we first checked if within each cluster there were 2 or more different 10x runs cells that had at least 100 cells within the cluster. If this condition was true, then these individual 10x runs cells were averaged from the log(GA + 1) matrix (maximum of 500 cells randomly sampled) to create pseudo-bulk replicates. If this condition did not satisfy for the cluster, and there was at least 1 10x run that had at least 100 cells and the remaining cells were at least 100 cells then the 1 10x run was averaged from the log(GA + 1) (maximum of 500 cells randomly sampled) to create a pseudo-bulk replicate. Then for the remaining cells a pseudo-bulk replicate was constructed by randomly sampling 100 cells from the remaining cells 250 times and we kept the sampling that produced the highest within cluster total log-variance (summed “rowVars” with the 1 10x run replicate). If that condition was not met then two replicates of 100 cells were randomly sampled 250 times and we kept the sampling that produced the highest within cluster total variance (summed “rowVars” with the both sampled replicates). Lastly if the cluster was smaller than 150 total cells the number of cells sampled was 2/3 the size of the cluster. This workflow was designed to construct pseudo-replicates from single-cell clusters that had high variance to attempt to capture real biological variation. In general, this approach is still an underestimate of variation when fully simulating replicates, so it’s important to be conservative when using these pseudo-replicates in further analyses.

#### Identification of cluster-specific peaks and gene activities through feature binarization

Once we had determined clusters from scATAC-seq data, we sought to identify peaks that were uniquely present within each cluster or combination of clusters. We modified a previously described approach to binarize each feature and then identify unique features in a simplistic manner^48^. For ATAC-seq, we used bulk pseudo-replicates and log-normalized the matrix using “edgeR::cpm(mat,log=TRUE,prior.count=3)”. For gene activities, we used bulk pseudo-replicates and re-normalized the matrix by first converting to gene activities and then converting back to log by computing “log2(edgeR::cpm(2^logMat−1)+1)”. Next we computed the mean and sd for each cluster using “rowMeans” and “rowSds” respectively in R. Next, for each feature peak or gene we ranked the clusters by their intra-cluster mean. Then, we iterated from the second lowest cluster asking whether the mean of that cluster: (1) is greater than the maximum intra-cluster mean plus the 0.5 times the intra-cluster standard deviation of the next-lowest cluster (1 sd for sub-cluster binarization), and (2): is greater than log2FC to the maximum intra-cluster mean of 0.25. This process was continued and the last time this criterion was met, the break point is labeled and all clusters above this intra-cluster mean were marked with a “1” and below a “0”. If a peak does not have a break point, it is discarded. This binarization will capture peaks that are unique to multiple groups. Next, all classified peaks/genes that correspond to more than a total of the floor of a third the clusters used as input. Next, for each peak/gene a t-test was computed comparing all “1’s” and “0’s” and the p-values were adjust for multiple hypothesis through the Benjamini-Hochberg correction by “p.adjust(method=“fdr”)”. All peaks/genes that had an adjusted p-value below 0.01 were kept. Lastly, we filtered out all binarization patterns that were classified less than 25 times. These were then plotted in R using the package ComplexHeatmap.

#### Pseudotime Analysis

To order cells in pseudotime, we sought to create a trajectory that we could align cells to. We chose to use UMAP for alignment if cells were part of a continuous substructure, since local distances are better preserved. First, we described candidate trajectories by ordering clusters. Next, for each cluster we calculated the mean coordinates in both dimensions and filtered cells that were in the top 5% euclidean distance to the mean coordinates. Next, we computed the distance for each cell from clusteri to the mean coordinates of clusteri+1 along the trajectory. We then computed a pseudotime vector by calculating the quantiles for each cell by their distance to the next cluster and added the current iteration. This allowed us for each of the cells (not filtered), have a UMAP coordinate and a time component. Next, we fitted a continuous trajectory to both UMAP coordinates using “smooth.spline” with dof =250 and spar = 1. Then, we aligned all cells to the trajectory by their euclidean distance to the nearest point along the manifold. We then computed then scaled this alignment to 100 and used this as pseudotime for further analyses.

To further support longer trajectories in pseudotime, we wanted to test whether the trajectory is significant by its ordering. To test our trajectories, we took the latest cluster and then ranked the top 10,000 accessible peaks and computed the euclidean distance to all other clusters (logCPM). We then continued in this reverse trajectory and computed the distance to all other clusters that do not include the previous clusters for directionality. To determine the significance of the ordering, we then permuted the ordering of the trajectory 5000 times and computed the average rank of the ordering for the permuted and input trajectory. This allowed for an empirical p-value calculation that we can then assign signified to this reduced dimension trajectory from the original accessibility matrix.

We then sought to create matrices that convey the pseudotime trends across features. To do this analysis, we ordered the cells by their pseudotime and fit a smoothed line for each feature by using “geom_smooth” with method “gam” and formula “y ~ s(x, bs = “cs”)” and n = 100. For peaks, this was done on the binarized sparse accessibility matrix, gene activities done using the log-normalized (log(GA + 1)) matrix, and chromVAR deviation scores were used in pseudotime analyses. We wanted to de-duplicate chromVAR CIS-BP motifs by correlating the gene activities of a TF to the inferred activity of chromVAR. We correlated TFs and their corresponding gene activities and then using cor.test in R kept the associated p-value and adjusted for multiple hypothesis through the Benjamini-Hochberg correction by “p.adjust(method=“fdr”)”. Next, we computed the quantile for each TFs gene activity average and variance. We then averaged these quantiles to equally weight the log-average gene activity and log-variance. We then filtered the top 25% of TFs by this criterion and then further by TF-motif pairs that were correlated above 0.35 and FDR < 0.001. These were then used to represent TF-motif pairs identified are more likely to be involved in gene regulation across the identified pseudotime.

#### Barnyard Mixing Analysis

We sought to assess the rate at which multiplets (more than one cell per droplet) occur at different cell loadings to estimate the relative multiplets in our data. To calculate this rate, we performed human (GM12878) and mouse (A20) mixing experiments at loadings of 500, 1,000, 5,000 and 10,000 cells. We aligned them using 10x Cell Ranger. We then filtered cells as described above for both hg19 and mm10 genomes (TSS enrichment of 8 and 1,000 unique fragments). Next, we determined the effective multiplet rate first by computing the fragments to each genome to the combined fragments and then labeled multiplets by assigning cells that have fractional fragments less than 0.95 to either hg19 or mm10.

We then wanted to determine the effect of different numbers of cells and unique fragments used for peak calling by down sampling cells and unique fragments. We did this analysis by merging all GM12878 and A20 identified fragments from the mixing experiments into 1 fragments file. Next, we down sampled the fragments file by first the number of cells and then the number of fragments to make the unique fragments per cell match the desired output. We then called peaks on each of these fragments by creating a bed file of the Tn5 insertions (ends of the fragments) with MACS2 callpeak command with parameters “--shift -75 --extsize 150 --nomodel --call-summits --nolambda --keep-dup all -q 0.05.” The peak summits were then extended by 250bp on either side to a final width of 501bp, filtered by the ENCODE hg19 blacklist (https://www.encodeproject.org/annotations/ENCSR636HFF/), and then filtered to remove peaks that extend beyond the ends of chromosomes. We then took the top 100,000 non-overlapping extended summits, as previously described^48^. We repeated this on the total fragments file to get a GM12878 and A20 peak set. Next then computed the fractional of peaks recovered by using “countOverlaps” and dividing by the extended summits of the total fragments file fin R. Lastly, we then counted the number of Tn5 insertions for each down sampled fragment file within the GM12878 and A20 peak set, log-normalized the matrix with “edgeR::cpm(mat, log = TRUE, prior.count = 3)” and computed the pearson correlation. We then plotted the results in R using “ggplot”.

#### Frozen vs Fresh Analysis

We wanted to compare the effect of sample preparation on scATAC-seq data quality. Thus, we performed scATAC-seq on PBMCs that were freshly isolated, frozen, and frozen sorted for live cells. We then filtered the cells as described above (TSS enrichment 8 and 1,000 unique fragments) and then used our TF-IDF cluster peak calling framework as described above to get a peak set for each experiment. We then subsequently used our TF-IDF cluster analysis as described above (top 25,000 variable peaks with SVD dimensions 1-50) and computed clusters for each experiment. We then computed the Receiver Operating Characteristic (ROC) curve for both frozen samples against the fresh sample by using “overlapsAny” and ranking the peaks by MACS2 score in R. We then computed PCA with “prcomp” and correlations between the identified clusters and entire experiment within the fresh samples ATAC-seq peaks.

#### Spike-Ins Analysis

We sought to test our sensitivity with our analysis workflow by performing scATAC-seq on monocyte and T cell mixtures at various loadings. We then filtered the cells as described above (TSS enrichment 8 and 1,000 unique fragments) and then used our TF-IDF cluster peak calling framework as described above to get a peak set for each experiment. We then subsequently used our TF-IDF cluster analysis as described above (top 25,000 variable peaks with SVD dimensions 1-50) and computed clusters for each experiment. We then used “RunUMAP” with default parameters from Seurat (v2.3) to compute a UMAP for each spike-in experiment. We then computed the gene activities scores as described above by using the full hematopoiesis peak set, accessibility matrix, and co-accessible links and added each sample individually for calculating the gene activity scores. We then computed a monocyte score by taking the log(GA+1) average for *CD14*, *MAFB*, *HLA-DRB1*, *TREML4*, *CSF1R*, *CEBPA*, *TLR4*, *HLA-DRA*, and *CD74*. Lastly, we computed a T cell score by taking the log(GA+1) average for *CD3E*, *CD2*, *CD5*, *CD7*, *IL7R*, *IL2*, *TCF7*, *CD3D*, and *CD3G*.

## Data Availability

All single-cell sequencing data have been deposited in the Gene Expression Omnibus (GEO) and are awaiting accessioning.

## Supplementary Figure Legends

**Supplementary Figure 1. Droplet-based scATAC-seq workflow and quality control measurements.** (**a**) Protocol steps for scATAC-seq in droplets. (**b**) Genome tracks showing the comparison of aggregate scATAC-seq profiles from A20 B lymphocytes (top panel). scATAC-seq profiles were obtained from four independent experiments, as indicated. The bottom panel shows accessibility profiles of 100 random single GM12878 cells from two cell mixing experiments. Each pixel represents a 100bp region. (**c**) Pearson correlation heatmaps of log-normalized reads in bulk GM12878 Omni-ATAC-seq peaks in aggregate ATAC-seq profiles generated from varying numbers of single cells, or from published Omni-ATAC profiles^5^. (**d**) Peak recovery analysis with subsampled cells and unique fragments as determined by x-axis and colors respectively. scATAC-seq cells were subsampled to the indicated unique fragments, and the proportion of peaks recovered from the aggregate profile was calculated as a function of number of cells analyzed. (**e**) Pearson correlation analysis with subsampled cells and unique fragments as determined by x-axis and colors respectively. scATAC-seq cells were subsampled to the indicated unique fragments, and pearson correlation to the aggregate profile was calculated as a function of number of cells analyzed. **(f)** Analysis workflow for scATAC-seq data in this study.

**Supplementary Figure 2. scATAC-seq performance in frozen cells and synthetic cell mixtures.** (**a**) Synthetic immune cell mixture quality control experiments. Sorted human monocytes or T cells were mixed at the indicated ratio and analyzed with scATAC-seq. Plots show the UMAP of scATAC-seq profiles (top), and gene score activity for monocyte- or T cell-associated cardinal genes (see Methods) in each single cell (middle and bottom). Dashed circles indicate monocyte and T cell identity of single cells as determined by ATAC-seq profiles. Colors indicate cluster identity defined *de novo*. (**b**) Sorted human monocytes or T cells were mixed at the indicated ratio and analyzed with scATAC-seq and analyzed as described in (a). (**c**) Comparison of data quality from fresh and frozen PBMCs, and frozen PBMCs sorted for live cells. Representative ATAC-seq data quality control filters by sample source. Shown are the number of unique ATAC-seq nuclear fragments in each single cell (each dot) compared to TSS enrichment of all fragments in that cell. Dashed lines represent the filters for high-quality single-cell data (1,000 unique nuclear fragments and TSS score greater than or equal to 8). (**d**) One-to-one plots of log-normalized reads in aggregated scATAC-seq in profiles generated from the indicated cell source (fresh, frozen, or frozen sorted PBMCs). Peaks were defined in fresh samples. Numbers indicate Pearson correlation value. (**e**) ROC curve showing recovery of fresh PBMC peaks with frozen or frozen sorted cells. (**f**) PCA of scATAC-seq clusters identified in fresh, frozen, or frozen-sorted PBMCs in fresh PBMC peaks.

**Supplementary Figure 3. Sample descriptions and quality control of scATAC-seq hematopoiesis data.**(**a**) UMAP projection of 63,882 scATAC-seq profiles of bone marrow and peripheral blood immune cell types. Dots represent individual cells, and colors indicate the experimental source of each cluster, as labeled on the right of the plot (see Methods). (**b**) Bar plots indicate the number of scATAC-seq profiles obtained from each experimental source of cells (left), and the median number of unique nuclear fragments in single cells (right). (**c**) UMAP projection of 63,882 scATAC-seq profiles of bone marrow and peripheral blood immune cell types. Colors represent the log10 number of unique nuclear fragments per single cell. (**d**) Representative scATAC-seq data quality control filters by sample source. Shown are the number of unique ATAC-seq nuclear fragments in each single cell (each dot) compared to TSS enrichment of all fragments in that cell. Dashed lines represent the filters for high-quality single-cell data (1,000 unique nuclear fragments and TSS score greater than or equal to 8). (**e**) single-cell ATAC-seq data quality control filters in profiles generated using the C1 microfluidic system^11^ (Fluidigm; left) or sci-ATAC-seq^12^ (middle and right panels).

**Supplementary Figure 4. *Cis*-regulatory elements in hematopoiesis and co-accessibility validation.** (**a**) Genome tracks of aggregate scATAC-seq data, clustered as indicated in Figure 2b. Arrows indicate the position and distance (in kb) of intragenic or distal enhancers in each gene locus. (**b**) MetaV4C plot of H3K27ac HiChIP data demonstrating enhancer interaction signal (EIS) at Cicero-identified co-accessible *cis*-elements (linked to promoter elements). Each plot shows the aggregate HiChIP signal (from 3 T cell types) between two linked *cis*-elements identified in scATAC-seq data. The peak indicates an enrichment of enhancer interaction signal (EIS) at the linked peaks compared to surrounding genomic regions. Biased links are identified by differential peak analysis in the indicated scATAC-seq clusters. (**c**) Support for Cicero-identified co-accessible *cis*-elements by GTEX eQTL data. Shown is the enrichment of eQTL signal (determined in the indicated tissue type) in co-accessible sites linked to promoter elements described in GTEX vs 250 permutations of ATAC-seq peak to genes. (**d**) Heatmap of log-normalized gene activity scores for the indicated genes. (**e**) UMAP projection colored by log normalized gene activity scores demonstrating the accessibility of *cis*-regulatory elements linked (using Cicero) to the indicated gene.

**Supplementary Figure 5. TF motif accessibility in hematopoiesis and cell type-specific GWAS enrichment.** (**a**) Example TF footprints and motifs in the indicated scATAC-seq clusters identified in Fig 2b. The Tn5 insertion bias track is shown below. (**e**) UMAP projection of scATAC-seq profiles colored by chromVAR TF motif bias-corrected deviations for the indicated factors. (**c**) Analysis workflow for GWAS enrichment scores using cicero co-accessibility. (**d**) Heatmap showing GWAS deviation scores for PICS SNPs associated with the indicated diseases. PICS SNPs were identified previously^22^. Example of increased ATAC-seq signal in a GWAS-containing cis-element in NK and T cell scATAC-seq clusters. The HiChIP plot (top) demonstrates increased Enhancer Interaction Signal (EIS) between the *STAT4* promoter and the highlighted *cis*-elements.

**Supplementary Figure 6. Sample descriptions and quality control of scATAC-seq profiles of the BCC TME.** (**a**) UMAP projection of 37,818 scATAC-seq profiles of BCC TME cell types. Dots represent individual cells, and colors indicate the experimental source of each cluster, as labeled on the right of the plot (see Methods). ‘Total’ samples were sorted as all live cells in a single BCC biospy. ‘T cell’ samples were sorted as CD3^+^ cells in the biopsy. ‘Immune’ samples were sorted as CD45^+^CD3^−^ cells in the biopsy. ‘Stromal’ samples were sorted as CD45^−^CD3^−^ cells in the biopsy. (**b**) Bar plots indicate the number of scATAC-seq profiles obtained from each experimental source of cells (left), and the median number of unique nuclear fragments in single cells (right). (**c**) UMAP projection of 37,818 scATAC-seq TME profiles. Colors represent the log10 number of unique nuclear reads per single cell. (**d**) Representative ATAC-seq data quality control filters by sample source. Shown are the number of unique ATAC-seq nuclear fragments in each single cell (each dot) compared to TSS enrichment of all fragments in that cell. Dashed lines represent the filters for high-quality single-cell data (1,000 unique nuclear fragments and TSS score greater than or equal to 8). (**e**) Genome tracks of aggregate scATAC-seq data, clustered as indicated in Figure 4b. Arrows indicate the position and distance (in kb) of intragenic or distal enhancers in each gene locus.

**Supplementary Figure 7. TF motif accessibility in the BCC TME.**(**a**) Example TF footprints and motifs in the indicated scATAC-seq clusters identified in Fig 4b. The Tn5 insertion bias track is shown below. (**e**) UMAP projection of scATAC-seq profiles colored by chromVAR TF motif bias-corrected deviations for the indicated factors.

**Supplementary Figure 8. Regulatory landscapes of tumor-infiltrating T cell subsets.** (**a**) UMAP projection of intratumoral T cell scATAC-seq data colored by log normalized gene activity scores, demonstrating the accessibility of *cis*-regulatory elements linked (using Cicero) to the indicated CD8^+^ T cell signature genes. (**b**) UMAP projection of intratumoral T cell scATAC-seq data colored by log normalized gene activity scores, demonstrating the accessibility of *cis*-regulatory elements linked (using Cicero) to the indicated CD4^+^ T cell signature genes. (**c**) Genome tracks of Tfh signature genes in aggregate scATAC-seq data, clustered as indicated in Figure 5a. Arrows indicate the position and distance (in kb) of intragenic or distal enhancers in each gene locus. (**d**) Heatmap of Z-scores of 35,147 *cis*-regulatory elements in scATAC-seq clusters derived from (b). Labels indicate cell type-specific accessibility of regulatory elements. (**e**) Genome tracks of CD8^+^ TEx signature genes in aggregate scATAC-seq data, demonstrating the overlap of TEx and Tfh regulatory elements. Arrows indicate the position and distance (in kb) of intragenic or distal enhancers in each gene locus. (**f**) Heatmap representation of ATAC-seq chromVAR bias-corrected deviations in the 250 most variable TFs across all intratumoral T cell scATAC-seq clusters, as identified in Figure 5a. Cluster identities are indicated at the bottom of the plot.

## Supplementary Table Legends

**Supplementary Table 1. Quality control sample characteristics.** Shown are the source and single cell numbers for each sample analyzed in QC experiments. Related to Fig. 1 and Supplementary Figs. 1-2.

**Supplementary Table 2. Hematopoiesis sample characteristics.** Shown are the source, isolation method, and single cell numbers for each sample analyzed by scATAC-seq in the study of immune cell development. Related to Figs. 2-3 and Supplementary Figs. 3-5.

**Supplementary Table 3. BCC sample characteristics.** Shown are the patient and cell characteristics for all BCC scATAC-seq samples. Response to anti-PD-1 therapy was categorized using RECIST scoring, as described in the Methods. Related to Figs. 4-5 and Supplementary Figs. 6-8.

