## Supplementary Figures for "Massively parallel single-cell chromatin landscapes of human immune cell development and intratumoral T cell exhaustion"

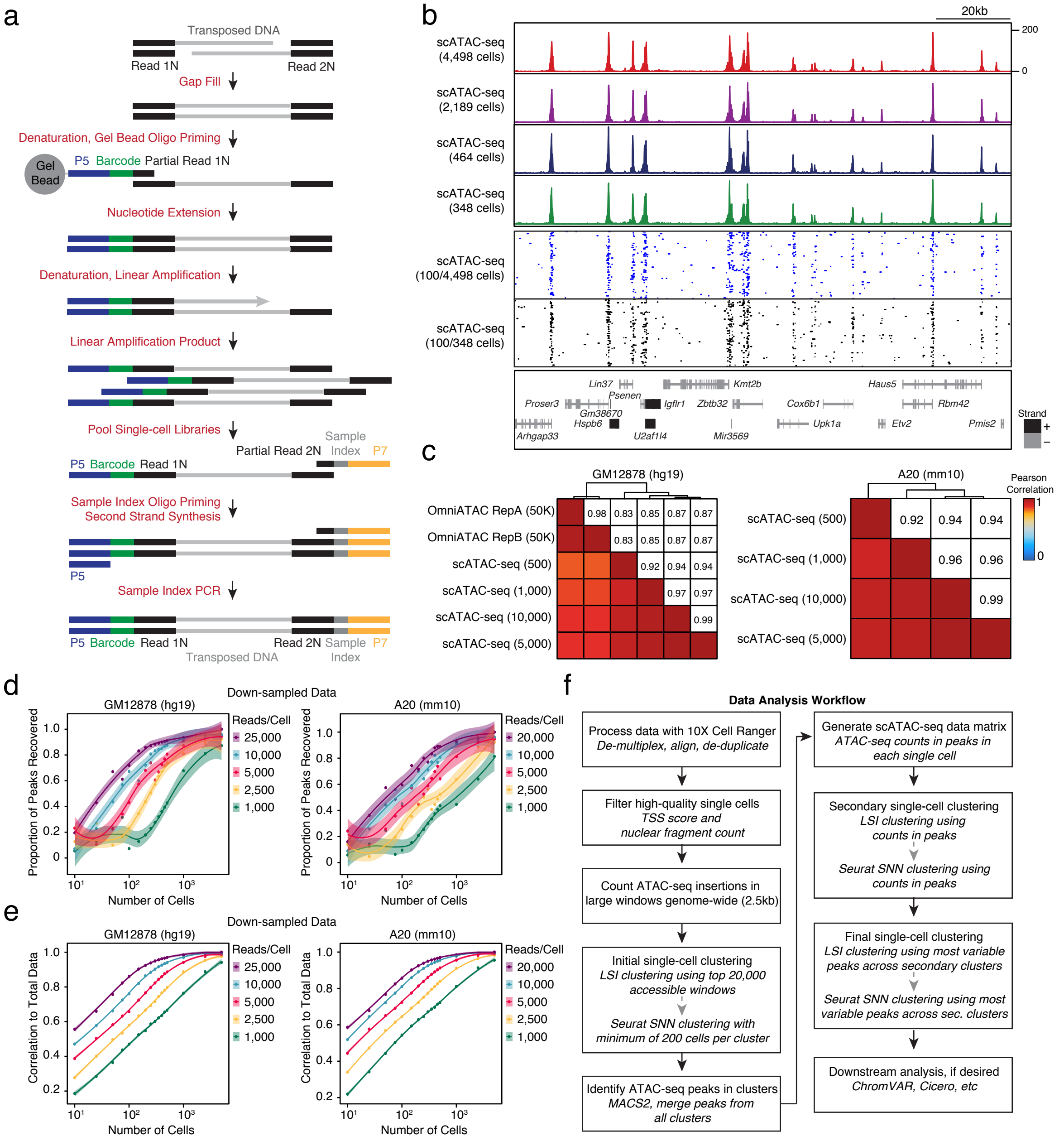


Supplementary Figure 1

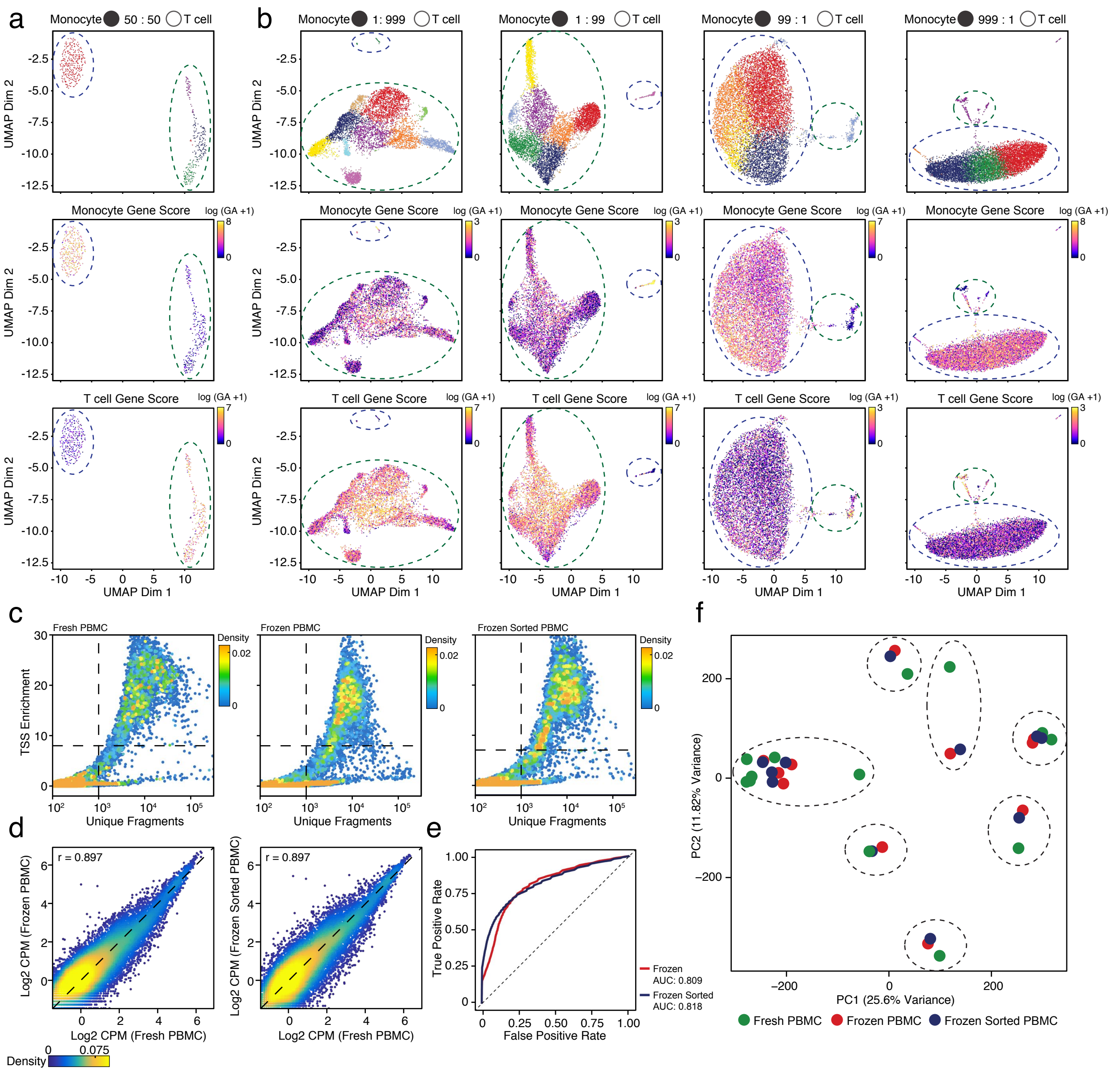


Supplementary Figure 2

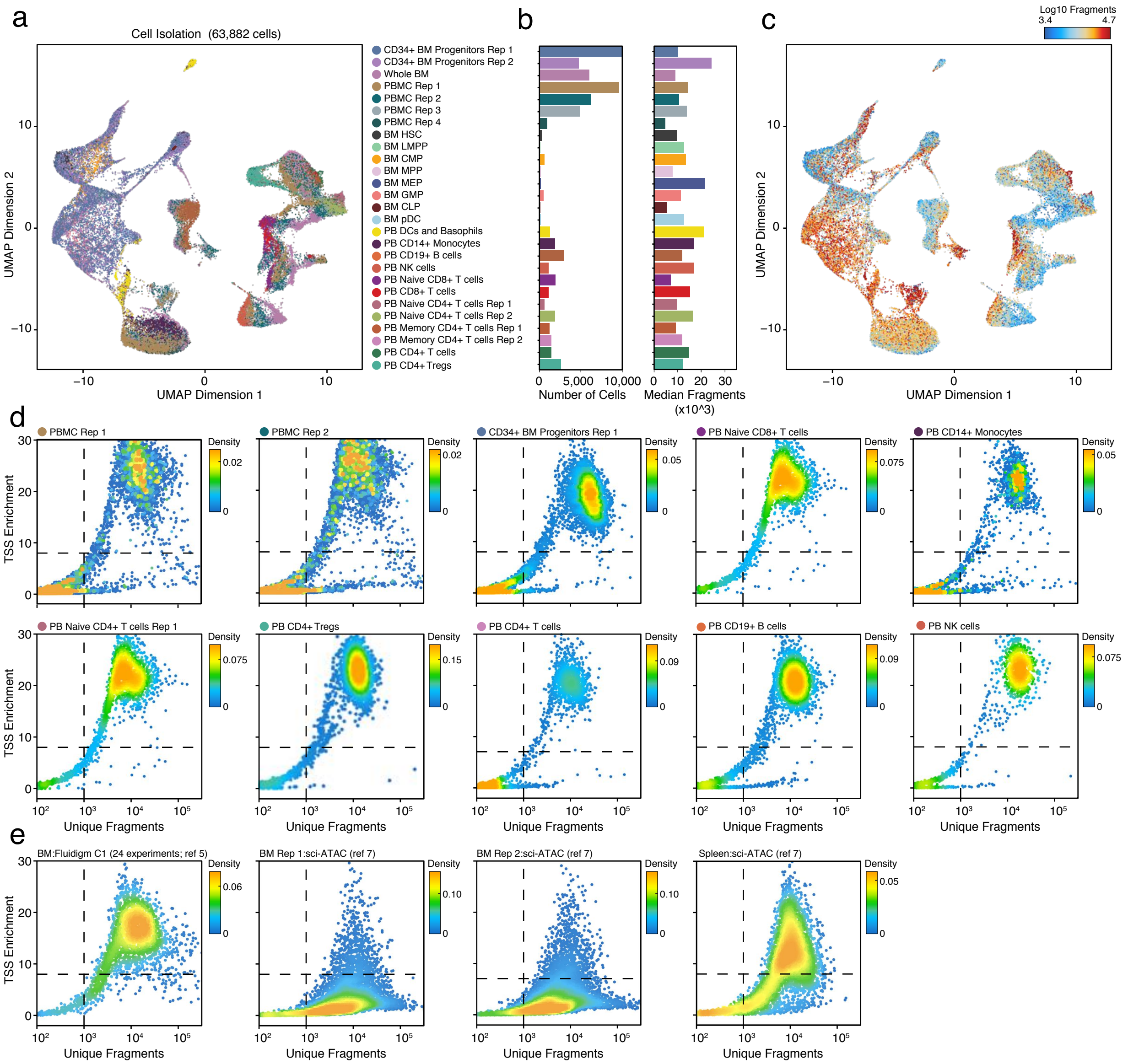


Supplementary Figure 3

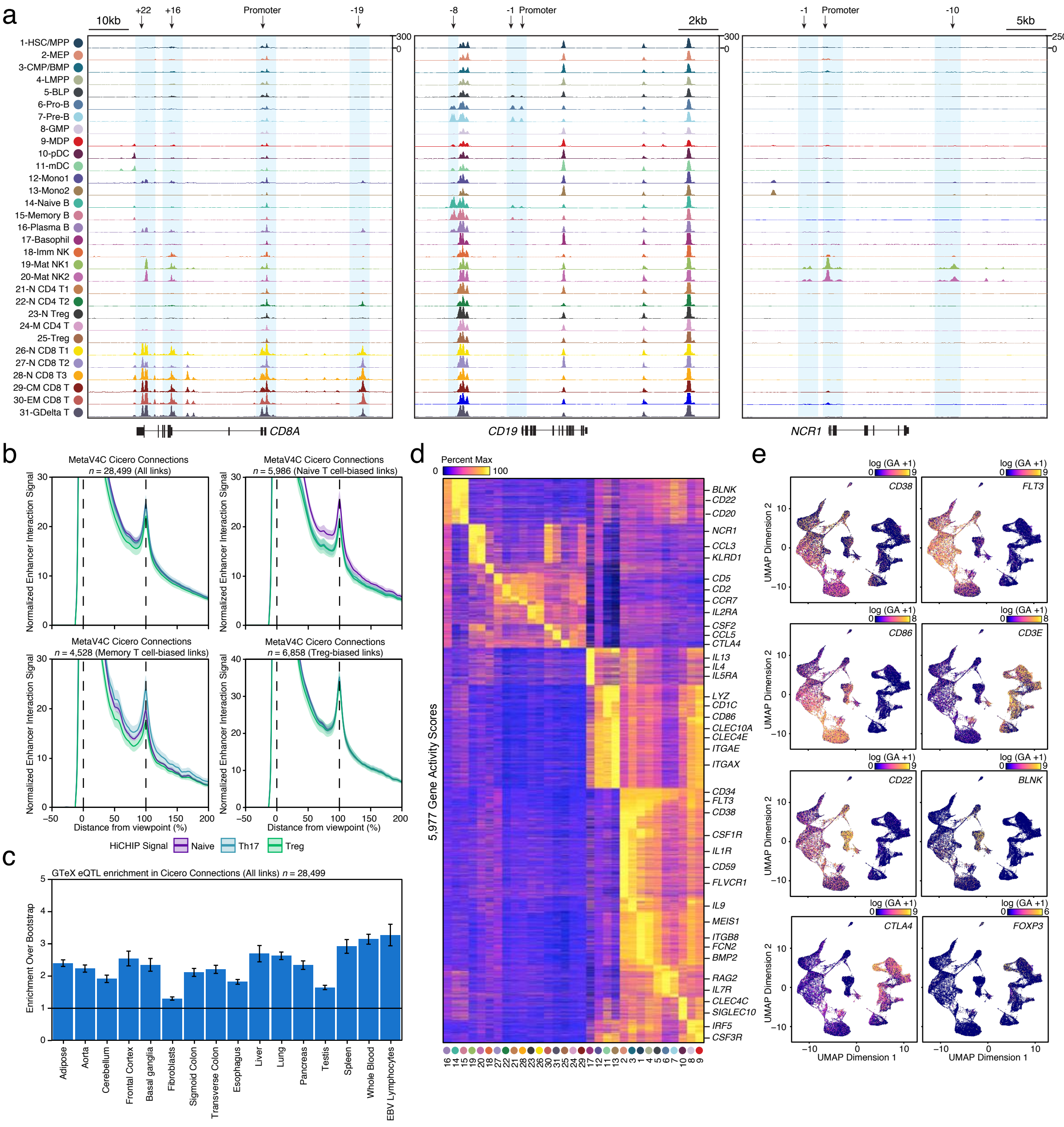


Supplementary Figure 4

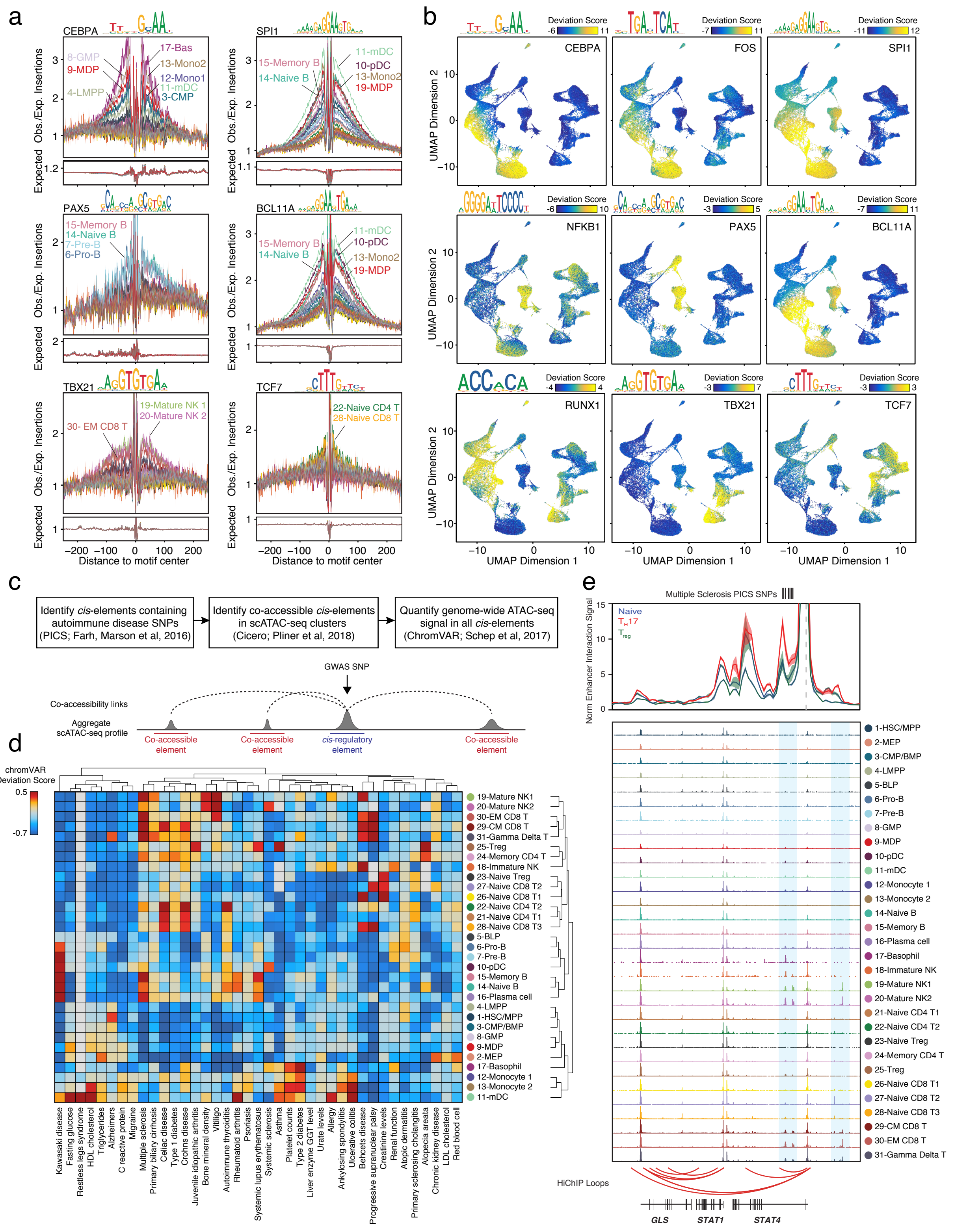


Supplementary Figure 5

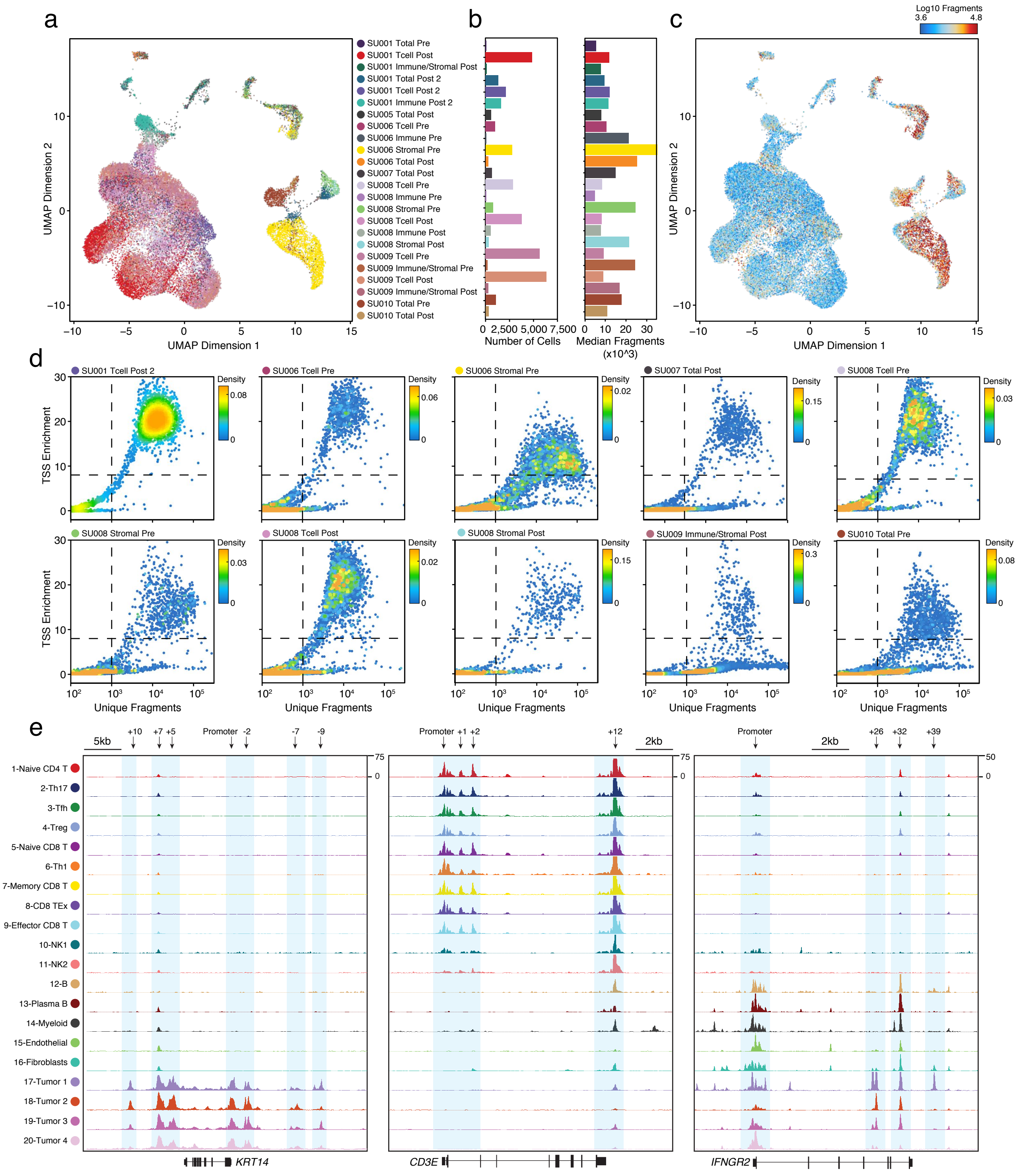


Supplementary Figure 6

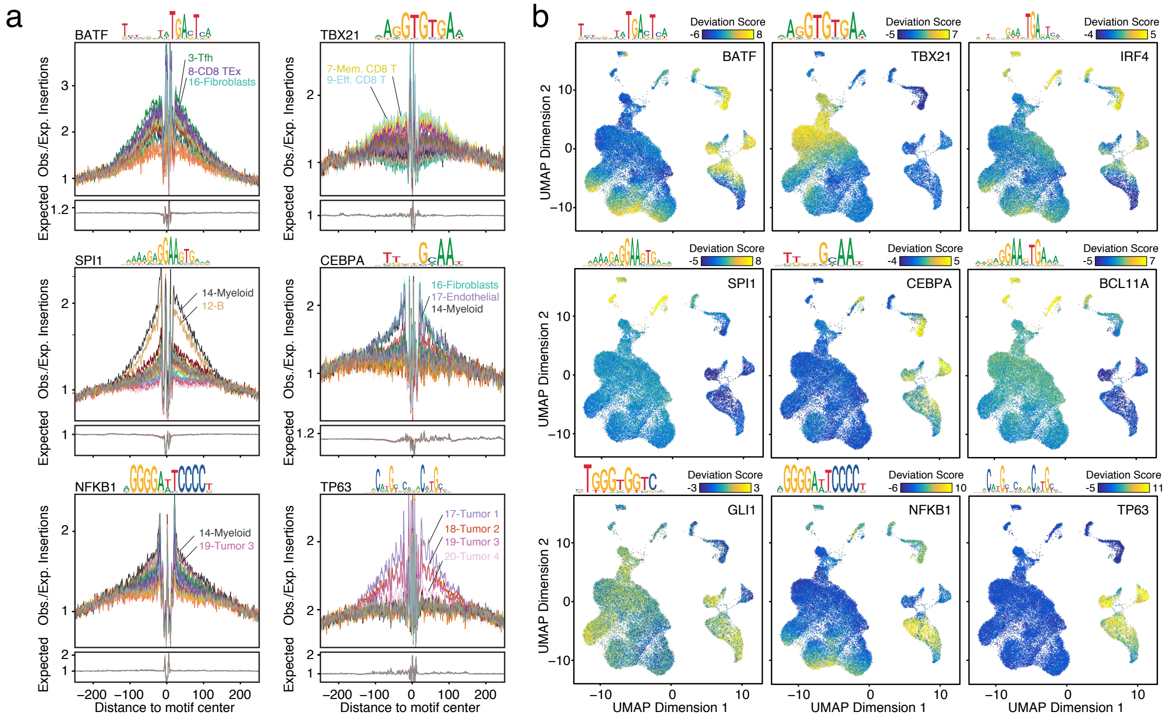


Supplementary Figure 7

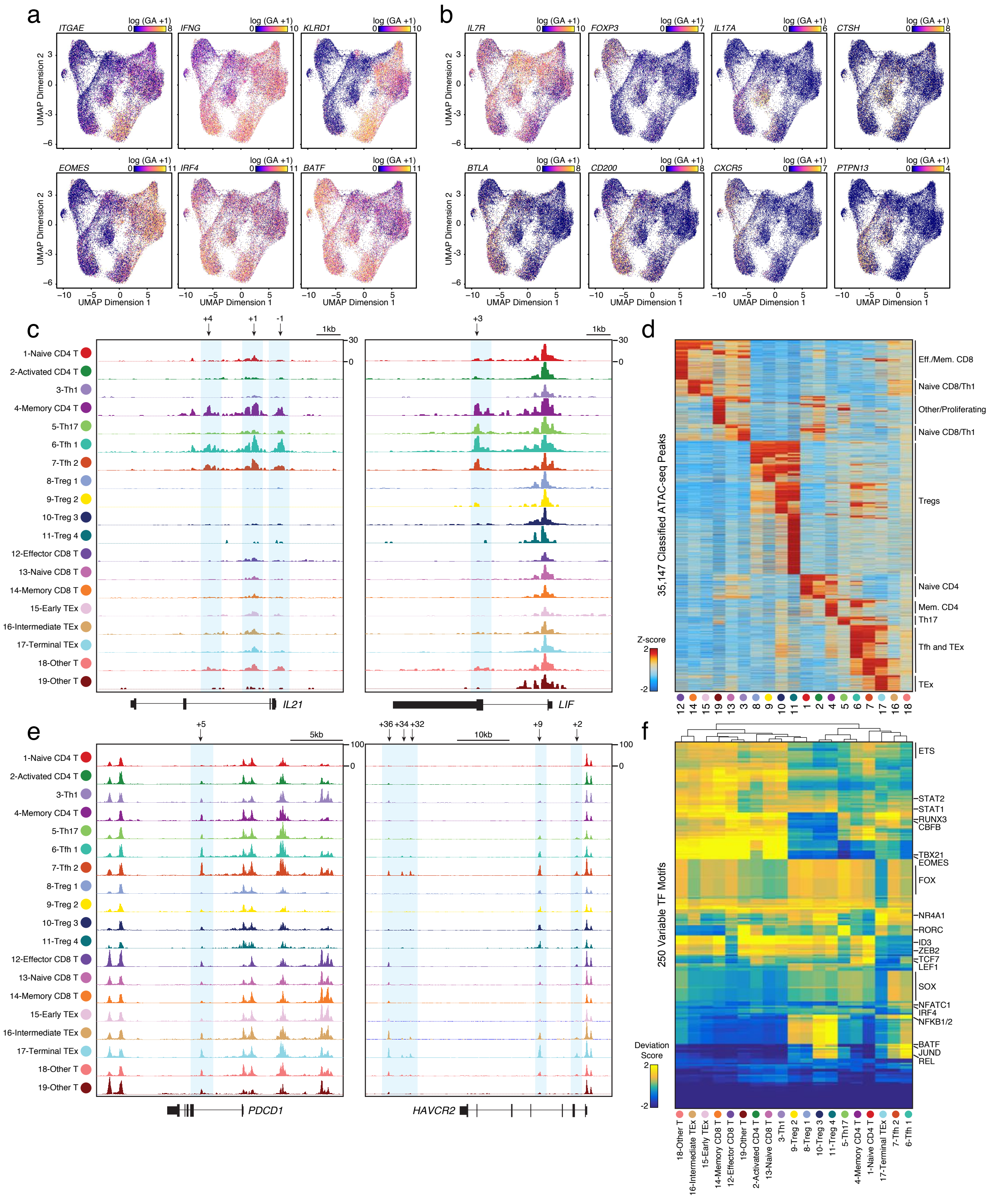


Supplementary Figure 8
